# Elp3 uses a conserved molecular tunnel to transport acetate between distant active sites and catalyze tRNA wobble base modification

**DOI:** 10.1101/2025.05.07.652618

**Authors:** Evan P. Geissler, Youmna Moawad, Paige N. Roehling, Katherine Martin, Papa Nii Asare-Okai, Jeffrey S. Mugridge

## Abstract

The radical SAM enzyme Elp3 and eukaryotic Elongator complex catalyze formation of a key intermediate transfer RNA (tRNA) modification, 5-carboxymethyluridine (cm^5^U), in the anticodons of tRNAs across all domains of life. cm^5^U-derived modifications are important for fine tuning codon-anticodon interactions and efficient protein translation, and defects in this modification are linked to development of neurodegenerative disease in humans. Here we reconstitute tRNA modification activity with a model Elp3 enzyme and combine structural analyses, enzymology, and isotope incorporation experiments to show Elp3 harbors a conserved molecular tunnel that shuttles free acetate molecules from the acetyl-CoA binding domain to the radical SAM active site over 20 Å away, where acetate undergoes radical-mediated reaction and addition to tRNA U34. Our model explains how Elp3 and Elongator bridge a large distance between active sites to catalyze tRNA carboxymethylation and illustrate a unique mechanism for intermediate transport in radical SAM enzymes.

**Graphical Abstract:** The radical SAM enzyme Elp3 installs a critical tRNA wobble base modification in organisms across all domains of life. Here, the authors show how Elp3 uses a conserved molecular tunnel to transport acetate between distant Elp3 active sites to catalyze tRNA carboxymethylation, revealing a new mechanism for Elp3 and Elongator-mediated tRNA modification and the first example of acetate transport through an enzyme tunnel.

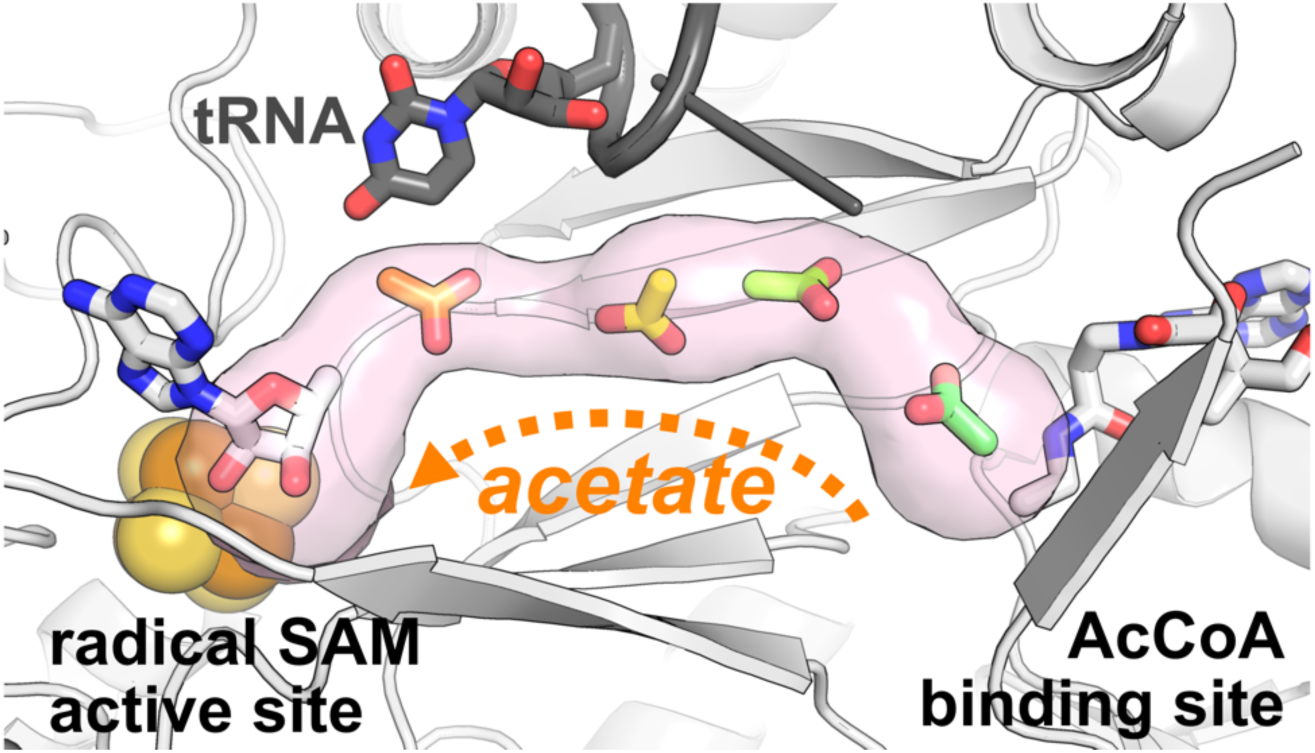

## Introduction

Organisms across all domains of life decorate their transfer RNA (tRNA) molecules with a wide variety of chemical modifications that finely tune protein translation and are critical for protein homeostasis and cellular health.^1–4^ Over 100 distinct tRNA nucleobase modifications have been identified to date,^5^ with an average of 13 modifications per tRNA found in human cells.^6^ 5-carboxymethyluridine (cm^5^U)-derived modifications (**Figure S1**) are found on over 50% of wobble uridines at position 34 in eukaryotic tRNAs^7^ and critically modulate the structure and function of the tRNA anticodon loop.^8^ This family of cm^5^U34- derived tRNA modifications facilitate base pair recognition between the mRNA codon and tRNA anticodon in the ribosome, ensuring efficient and accurate decoding and protein translation.^9–11^ Defects in the cm^5^U34-derived modification pathways lead to detrimental cellular phenotypes and disease outcomes. For example, proper levels of 5- methoxycarbonylmethyluridine 34 (mcm^5^U34)-modified tRNAs are required for effective translation of multiple DNA damage repair proteins and correct progression through the cell cycle in yeast,^12–14^ while reduction of mcm^5^U34 levels impairs selenocysteine protein expression and increases cellular sensitivity to reactive oxygen species in mammals.^15–17^ Loss of the related 5-methoxycarbonylmethyl-2-thiouridine (mcm^5^s^2^U) modification leads to frameshift errors during translation^18^ and dysregulation of metabolic^19^ and protein homeostasis^20–22^ in yeast, and is strongly linked to the development of neurodegenerative diseases and neurodevelopmental disorders, including familial dysautonomia, in humans.^23^

In eukaryotes, the multiprotein Elongator complex is responsible for installing cm^5^U34 on tRNA and additional tRNA-modifying enzymes subsequently act on cm^5^U34 to generate the family of cm^5^U34-derived modifications described above (**Figure S1**).^15,24–28^ The Elongator complex is composed of two copies each of Elongator proteins 1-6 (Elp1-6), where Elp3 is the enzymatic core of the complex that carries out the tRNA modification reaction and the other Elp proteins are thought to act as adaptor or scaffolding proteins that facilitate tRNA binding and release.^29,30^ In most archaea and numerous bacteria, only the catalytic subunit Elp3 is conserved and so this enzyme likely performs the tRNA modification reaction alone in those organisms where cm^5^U is formed.^31^ Structures of Elp3 have been determined from all three domains of life, revealing a highly conserved architecture consisting of a radical *S*-adenosylmethionine (rSAM) domain that binds a [4Fe-4S] cluster and cofactor SAM, and a lysine acetyltransferase (KAT) domain that binds acetyl-CoA (AcCoA) (**Figure S2**).^32–35^ Mutations in Elp3 and Elongator disrupt protein folding and stress responses,^36,37^ and are closely associated with the development and progression of a variety of human diseases, including various cancers^38–41^ and neurodevelopmental^42,43^ and neurodegenerative disorders^11,44^ such as familial dysautonomia^45^ and amyotrophic lateral sclerosis (ALS).^46,47^

Elp3 is thought to use a multistep mechanism in which tRNA binding triggers canonical rSAM chemistry at the [4Fe-4S] cluster to reductively cleave SAM and generate a reactive 5′-deoxyadenosyl radical (5′-dA·), which subsequently reacts with AcCoA or the acetyl group from AcCoA, to ultimately generate the cm^5^U34 modification on tRNA.^31^ However, recent structures of Elp3 and the Elongator complex with bound tRNA all show that the [4Fe-4S] active site on the rSAM domain and the AcCoA binding site on the KAT domain are separated by over 20 Å.^30,32–35^ This raises critical questions about how SAM (or 5′- dA·) and AcCoA cofactors come together in the Elp3 rSAM active site to react with tRNA and form the key cm^5^U wobble base modification. To date, there has only been a single report from 2014 in which any Elp3- or Elongator-mediated tRNA modifying activity has been detected with recombinant enzyme(s) *in vitro*;^31^ this challenge has so far prevented careful enzymological investigation and clear understanding of how Elp3 and Elongator modify tRNA.

Here we successfully reconstitute the *in vitro* cm^5^U34 tRNA modification reaction for the first time in over a decade using a model, recombinant archaeal Elp3 and combine structural analysis with enzymology and isotope incorporation experiments to propose a new mechanism for Elp3-mediated tRNA modification. We show that Elp3-tRNA complexes contain a conserved, enclosed molecular tunnel connecting the distant rSAM and KAT sites, that acetate can be used in place of cofactor AcCoA to carry out the cm^5^U modification reaction, and that blocking the molecular tunnel disrupts Elp3-mediated tRNA modification activity without significantly affecting tRNA binding or AcCoA binding and turnover. Together these data suggest a new model for Elp3- and Elongator-mediated tRNA modification in which the acetyl group from AcCoA is delivered to the rSAM active site via transport of free acetate molecules through a conserved, 20+ Å-long molecular tunnel connecting both domains of Elp3. We propose that key, conserved residues in the rSAM active site then noncovalently coordinate and position acetate for sequential reaction with 5′-dA· and tRNA U34 to catalyze cm^5^U formation on tRNAs from archaea to humans. This novel tRNA modification mechanism suggests Elp3 might be therapeutically targeted by inhibitors mimicking acetate intermediates, reveals the first example of acetate transport between enzyme active sites, and shows how molecular tunnels may be formed by protein-nucleic acid interactions in rSAM enzymes.

## Results

### Elp3-tRNA complexes form a conserved molecular tunnel connecting the rSAM active site and AcCoA binding site

Elp3 uses cofactors SAM and AcCoA to catalyze formation of cm^5^U by radical-mediated incorporation of the acetyl group from AcCoA into tRNA. In their 2014 study, Selvadurai *et al.* carried out isotope incorporation experiments using d3-acetyl-CoA that showed the 5′-dA· generated in the rSAM site directly abstracts a H atom (or D atom in the case of d3-AcCoA) from AcCoA’s acetyl group.^31^ However, every structure of Elp3 alone or in the context of the Elongator complex determined to date shows a 20+ Å distance between the rSAM active site where 5′-dA· is generated and the KAT domain where AcCoA is bound,^30,32–35^ raising the critical question of how the acetyl group from AcCoA comes close enough to react with 5′-dA·. Using a previously determined cryo-EM structure of tRNA-bound Elp3 from the yeast Elongator complex (PDB 8ASW),^30^ we performed a structural analysis using CAVER 3.0, which maps and analyzes tunnels and channels in protein structures.^48^ Our analysis revealed that the Elp3-tRNA complex forms an enclosed molecular tunnel reaching from the AcCoA binding site on the KAT domain directly to the [4Fe-4S] cluster and SAM binding site in the rSAM domain (**Figure 1A**). Formed mostly by the rSAM domain’s partial TIM barrel motif, the tunnel has additional surfaces provided by both the KAT domain and the substrate tRNA. The tunnel-lining residues are highly conserved, composed almost entirely of either identical or chemically similar residues in Elp3 sequences across all domains of life (**Figure S3**). In the yeast Elp3-tRNA structure, the diameter of the tunnel ranges from 3.4 to 6.4 Å, suggesting the possibility that small molecules could diffuse or be transported along its length.

**Figure 1.**
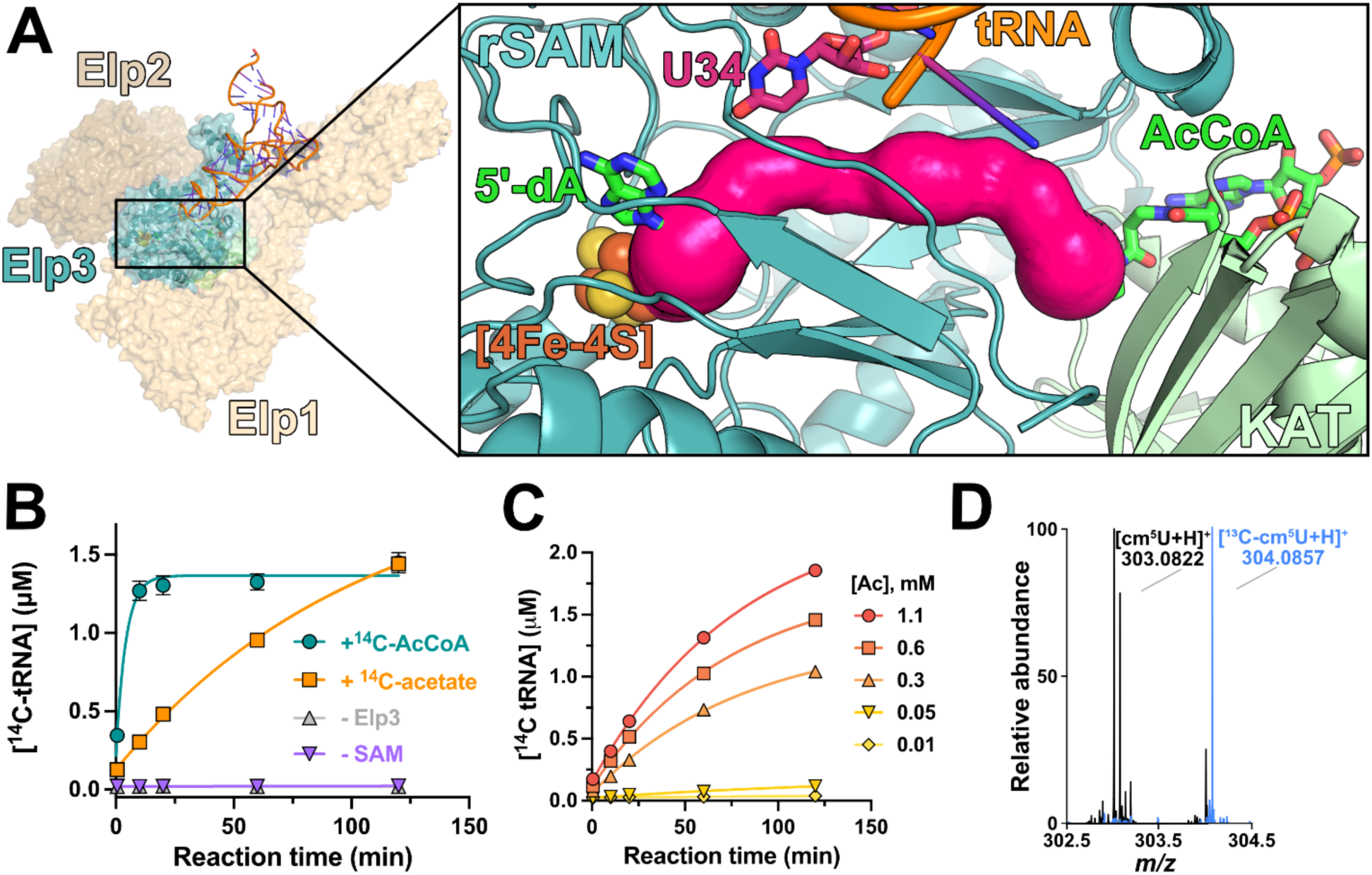
Elp3 uses a molecular tunnel to transport acetate to the rSAM active site for tRNA modification. **(A)** CAVER analysis of the yeast Elp1-Elp2-Elp3 complex bound to tRNA (PDB 8ASW) reveals an enclosed molecular tunnel connecting the Elp3 rSAM active site and the KAT domain AcCoA binding site (inset; CAVER-calculated tunnel shown in pink). The Elp3 rSAM domain is colored teal and the Elp3 KAT domain is colored light green; 5′-dA and AcCoA analog desulfo-CoA (aligned from PDB 6IA6) are shown as green sticks, but were not present in the CAVER calculation. tRNA and substrate tRNA base U34 are shown in orange and pink, respectively. **(B)** Upon incubation with substrate tRNA Arg^UCU^ and ^14^C-labeled acetate (orange squares), *Min* Elp3 carboxymethylates tRNA in similar quantities as the enzyme does with ^14^C- labeled AcCoA (teal circles) after 2 hours, while negative controls without Elp3 (gray triangles) or SAM (purple triangles) produce no radiolabeled tRNA product. Triplicate time courses were performed with 5 µM Elp3, 4.4 µM tRNA, 25 µM SAM, 0.5 mM dithionite as reductant, and either 27.5 µM ^14^C-AcCoA or 2 mM ^14^C-acetate, with data fit to a single-phase exponential equation and errors shown as SD. **(C)** Incubation of *Min* Elp3 with substrate tRNA Arg^UCU^ and increasing concentrations of ^14^C-acetate results in increasing amounts of ^14^C-labeled tRNA product formation. Time courses were performed with 5 µM Elp3, 4.4 µM tRNA, 25 µM SAM, 0.5 mM dithionite as reductant, and ^14^C-acetate concentrations as shown, with data fit to a single-phase exponential equation. **(D)** LC-MS analysis confirms that Elp3 forms ^12^C-containing (*i.e.* non- isotopically labeled) cm^5^U from ^12^C acetate *in vitro* (black spectrum), while an activity assay with ^13^C-labeled acetate forms ^13^C-labeled cm^5^U with an observed +1 Da mass shift in the cm^5^U nucleoside product (blue spectrum). tRNA modification assays for LC-MS analysis were performed with 5 µM Elp3, 4.4 µM tRNA, 25 µM SAM, 0.5 mM dithionite as reductant, and 10 mM ^12^C-(unlabeled) or ^13^C-acetate.

### Elp3 uses acetate as a key intermediate to form cm^5^U34

Based on our structural analysis with CAVER and discovery of the conserved molecular tunnel, we hypothesized that Elp3 might resolve the large distance between the [4Fe- 4S]/SAM and AcCoA binding sites by generating free acetate (or acetic acid, depending on local pH) from AcCoA at the KAT domain, transporting the acetate molecule through the tunnel, and delivering it for reaction with 5′-dA· and tRNA in the rSAM active site. To experimentally test this and other mechanistic hypotheses, we first overexpressed and anaerobically purified archaeal *Methanocaldococcus infernus* (*Min*) Elp3 and then screened [4Fe-4S] cluster reconstitution and reduction conditions in order to obtain *in vitro* tRNA-modifying activity. We verified the presence of [4Fe-4S] cluster in reconstituted Elp3 by UV-vis absorption at 420 nm (**Figure S4**) and then carried out radiolabel-based kinetic experiments using ^14^C-AcCoA to monitor Elp3-mediated tRNA modification over time. Under optimized anaerobic reaction conditions, we obtained robust and reproducible Elp3- and SAM-dependent tRNA modification activity and validated formation of cm^5^U on tRNA by LC-MS and MS/MS (**Figure S5**).

With catalytically active recombinant Elp3 in hand, we next tested if acetate was able to replace AcCoA in our ^14^C-based tRNA modification assays. Using ^14^C-labeled acetate in place of ^14^C-AcCoA, we found that Elp3 could still carry out the tRNA modification reaction (**Figure 1B**), and that the amount of ^14^C-tRNA product formed was dependent on the ^14^C- acetate concentration in the reaction (**Figure 1C**). We confirmed that cm^5^U was indeed being formed on tRNA during these reactions using unlabeled acetate and LC-MS (**Figure 1D**, black spectrum; **Figure S6**). Additionally, we carried out tRNA modification reactions using ^13^C-labeled acetate followed by LC-MS and observed a +1 Da mass unit shift in the resulting cm^5^U nucleoside product (**Figure 1D**, blue spectrum; **Figure S6**), which unambiguously demonstrates that the added acetate is the reactive species that Elp3 uses to modify tRNA in these assays. Together, these biochemical data show that Elp3 can entirely bypass the need for cofactor AcCoA in its tRNA modification reaction and instead use molecular acetate to catalyze cm^5^U formation.

### Blocking the molecular tunnel inactivates Elp3

If Elp3 uses a tunnel to transport molecular acetate from the KAT domain to the rSAM domain, sterically occluding the tunnel to restrict acetate transit should lead to diminished tRNA modification activity. Using the yeast Elp3-tRNA structure, we modeled how potential point mutations that increase the size of residues along the tunnel would narrow its diameter and identified the I154W mutation as one that would be predicted to substantially block the molecular tunnel (**Figure 2A**). Elp3 I154W was expressed, purified, and reconstituted using the same anaerobic conditions as for wild-type Elp3, resulting in purified recombinant enzyme that has the same size exclusion chromatography profile and UV-vis absorbance spectra as WT Elp3 (**Figure S7**). tRNA modification assays with ^14^C-AcCoA reveal that the I154W variant completely eliminates Elp3-mediated tRNA modification activity (**Figure 2B**). Critically, while I154W entirely abolishes tRNA modification activity, tRNA binding and AcCoA turnover are not significantly affected by this mutation compared to WT Elp3 (**Figure 2C & D**). These controls show that although I154W is located in relatively close proximity to the tRNA backbone in the Elp3-tRNA structure, the mutation does not impact tRNA binding or the binding and turnover of AcCoA, which is thought to be somehow coupled to formation of the Elp3-tRNA complex.^31^ These experiments suggest that blocking the molecular tunnel inactivates Elp3 tRNA modification catalysis by preventing passage of molecular acetate from the KAT domain into the rSAM active site after tRNA binding and AcCoA hydrolysis.

**Figure 2.**
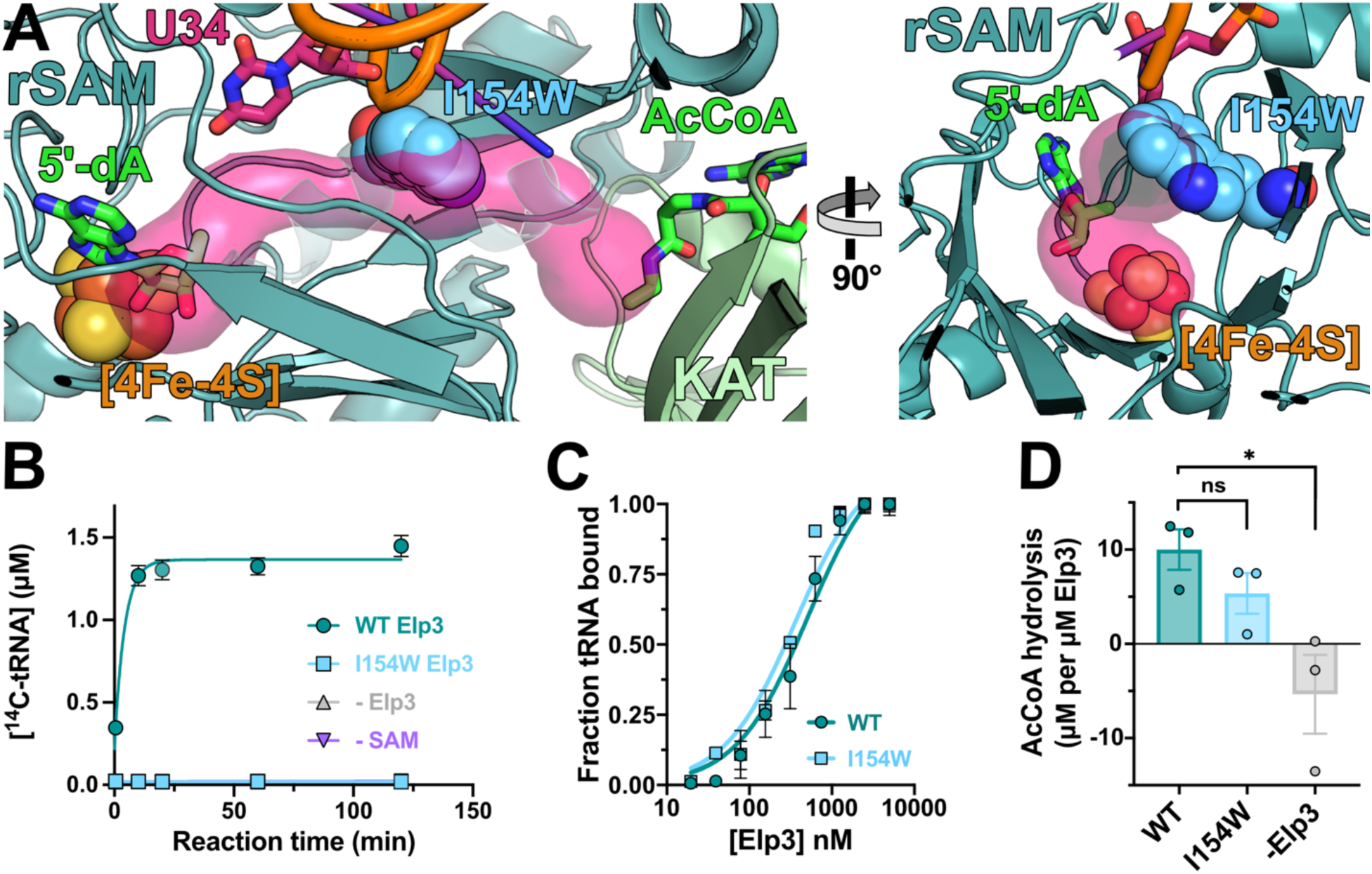
The Elp3 I154W mutation blocks the molecular tunnel and eliminates tRNA-modifying activity without disrupting tRNA binding or AcCoA hydrolysis. **(A)** Mutagenesis modeling and analysis of Elp3 bound to tRNA (from PDB 8ASW) in PyMOL suggested that the I154W mutation (shown in blue spheres), located midway along the molecular tunnel, will substantially block Elp3’s tunnel and likely impede diffusion of acetate from the KAT AcCoA binding site to the rSAM active site. **(B)** Incubation of Elp3 I154W with substrate tRNA Arg^UCU^ and ^14^C-labeled AcCoA *in vitro* results in no observable tRNA modification activity after 2 hours (light blue squares) compared to WT *Min* Elp3 (teal circles); activity of the I154W mutant is comparable to -Elp3 (gray triangles) and -SAM (purple triangles) negative controls. Triplicate time courses were performed with 5 µM Elp3, 4.4 µM tRNA, 25 µM SAM, 0.5 mM dithionite as reductant, and 27.5 µM ^14^C-AcCoA, with data fit to a single-phase exponential equation and errors shown as SD. **(C)** Elp3 I154W (light blue squares) binds substrate tRNA Arg^UCU^ with comparable affinity to WT Elp3 (teal circles). Elp3-tRNA binding was measured by triplicate electrophoretic mobility shift assays (EMSAs; **Figure S8**), where bound and unbound tRNA species at different Elp3 concentrations were separated, visualized by SYBR Gold staining, and unbound tRNA species were quantified using ImageJ to obtain fraction bound values. Data were fit to a single-site binding model with errors shown as SD. **(D)** Elp3 I154W (light blue) has similar AcCoA hydrolysis activity in the presence of substrate tRNA Arg^UCU^ compared to WT Elp3 (teal); both WT and I154W Elp3 show more AcCoA hydrolysis activity than a -Elp3 negative control (gray). AcCoA hydrolysis was measured in triplicate with a commercial fluorometric AcCoA quantification kit and statistical significance was calculated using a one-way ANOVA with Dunnett’s multiple comparisons test (* p<0.05, ns p>0.05); errors shown as SEM.

### Nucleophilic or hydrolytic reaction of acetate is not required for Elp3-mediated tRNA modification

Previously proposed mechanisms^30,31,35^ have suggested the possibility that acetylation of key Elp3 residues might be required to transport or help position the acetyl group in the rSAM active site for tRNA modification. To determine whether or not acetate forms an intermediate acetylated residue or undergoes hydrolysis during the course of the tRNA modification reaction, we performed an isotope incorporation experiment using doubly ^18^O-labeled acetate. If acetate must undergo nucleophilic addition and/or hydrolysis at any point during the reaction mechanism, this should result in the obligate loss of at least one of the two ^18^O isotope labels in the final cm^5^U product (**Figure 3A, i**). In contrast, if Elp3 uses only noncovalent coordination of free acetate to carry out the modification reaction, this would result in both ^18^O labels being retained in the final cm^5^U product (**Figure 3A, ii**). We carried out tRNA modification reactions using doubly ^18^O-labeled acetate and observed a peak corresponding to cm^5^U nucleoside product with a +4 Da mass unit shift, consistent with formation of cm^5^U that retains both ^18^O labels (**Figure 3B**, **Figure S9**). While peaks corresponding to non-labeled and singly ^18^O-labeled cm^5^U product are also present (**Figure S10**), this is likely due to oxygen’s known tendency to undergo exchange at carboxylate positions in water.^49,50^ The clear presence of doubly ^18^O-labeled cm^5^U product in Elp3 reactions carried out with ^18^O2-acetate strongly suggests that acetate is not required to undergo nucleophilic or hydrolytic reaction during the tRNA modification reaction cycle and is instead likely positioned in the rSAM active site through noncovalent interactions.

**Figure 3.**
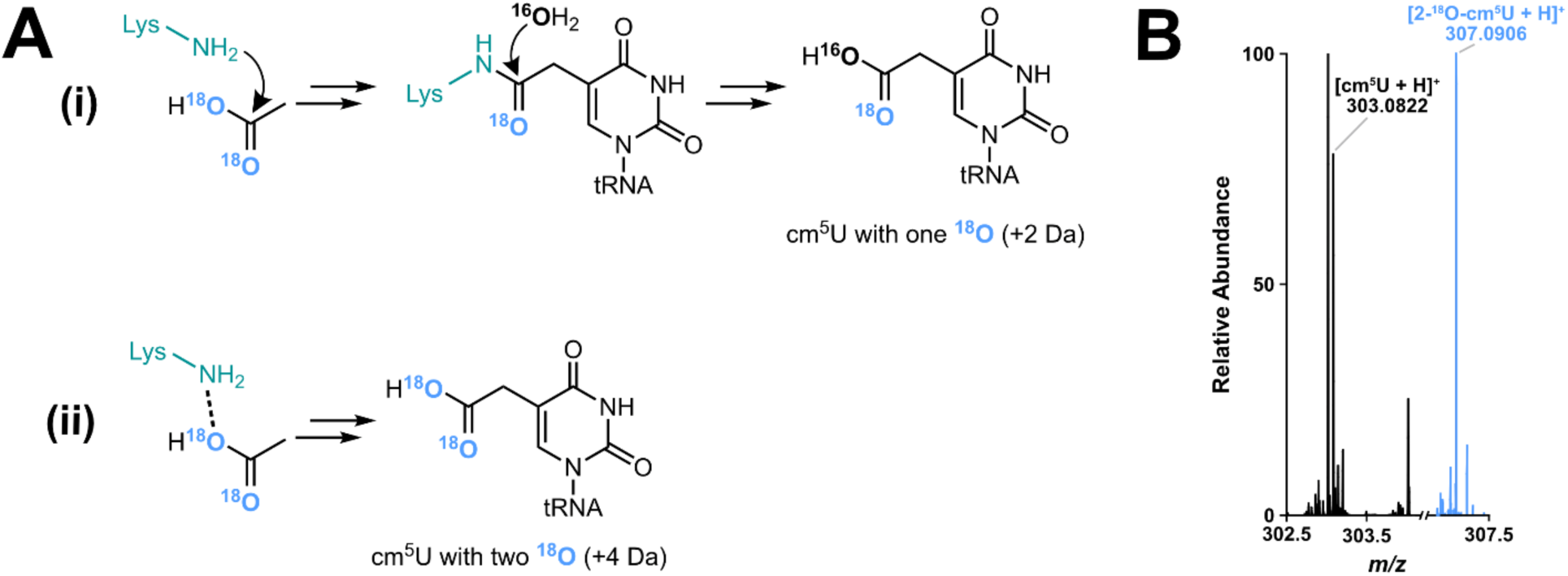
Elp3 forms doubly ^18^O-labeled cm^5^U tRNA from ^18^O2 acetate. (A) **(i)** If acetate undergoes nucleophilic attack during the Elp3-mediated reaction to form an acetylated residue, the reaction would displace one of the ^18^O isotope labels. Subsequent hydrolysis of the acetylated residue by bulk (^16^O, non-labeled) water would result in a cm^5^U product with one ^16^O and one ^18^O label in the carboxylate and a +2 Da mass unit shift above normal isotopic abundance cm^5^U. **(ii)** In contrast, noncovalent coordination of free acetate would result in both ^18^O labels being retained at the carboxylate and a +4 Da mass unit shift above normal isotopic abundance cm^5^U. **(B)** LC-MS analysis confirms that Elp3 forms doubly ^18^O-labeled cm^5^U from ^18^O2-acetate *in vitro*, with the resulting cm^5^U product (blue spectrum) showing a +4 Da mass shift compared to cm^5^U formed using normal isotopic abundance acetate (black spectrum). The normal abundance activity assay was performed with 5 µM Elp3, 4.4 µM tRNA, 25 µM SAM, and 10 mM normal isotopic abundance acetate and the ^18^O-acetate activity assay was performed with 25 µM Elp3, 4.4 µM tRNA Arg^UCU^, 50 µM SAM, and 10 mM ^18^O2-acetate.

### Conserved Elp3 rSAM active site residues play a key role in noncovalent positioning of acetate for radical-mediated reaction

Based on the ^18^O labeling experiments above, we presumed that a number of conserved rSAM active site residues of Elp3 would be used to noncovalently coordinate and position acetate for H atom abstraction by 5′-dA· and subsequent addition to tRNA U34. Although we have not yet succeeded in obtaining acetate-bound Elp3 structures, we used the molecular docking programs CaverDock^51^ and SeamDock^52^ to visualize and generate hypotheses about how acetate might be bound at different locations along the Elp3 molecular tunnel and within the rSAM active site (**Figure S11**). From these computational tools, we obtained predicted poses of acetate in the rSAM active site in which the acetate oxygen atoms hydrogen bond and make ionic interactions with several strictly conserved Elp3 residues (*Min* residues E207, K302, Y304; see **Table S1** for eukaryotic residue numbering), and at the same time position the acetate methyl group between the 5′ carbon atom of 5′-dA and the C5 position of U34 tRNA, which would allow direct H atom abstraction and radical transfer from 5′-dA· to the acetate methyl group and then to tRNA U34 (**Figure 4A**). While these are only rough models based on simple docking algorithms, the predicted acetate poses suggest that acetate can reasonably be accommodated at many positions along the molecular tunnel and oriented within the active site by key noncovalent interactions to facilitate sequential reaction with 5′-dA· and tRNA U34.

**Figure 4.**
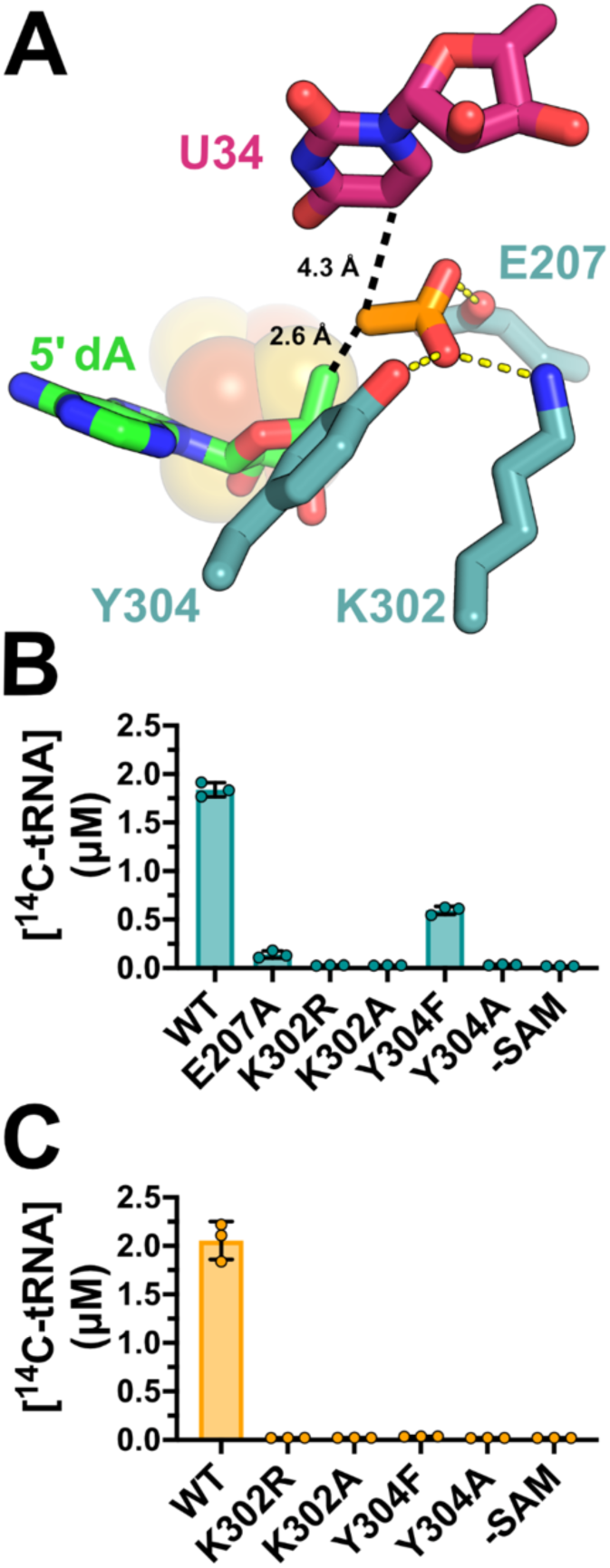
Molecular docking and activity assays reveal rSAM active site residues critical for Elp3- mediated tRNA-modifying activity. **(A)** SeamDock model of acetate bound in Elp3’s rSAM active site using the yeast Elp3-tRNA structure (PDB 8ASW), with *Min* Elp3 numbering. This model predicts hydrogen bonding and/or ionic contacts between acetate and conserved rSAM site residues E207, K302, and Y304. The Elp3 rSAM domain residues are colored teal, 5′-dA is shown in green, substrate tRNA base U34 is pink, and the [4Fe-4S] cluster is shown as spheres. Yellow dotted lines denote hydrogen bond or ionic interactions between 2.9 and 3.2 Å; black dotted lines show the distances between the acetate methyl carbon and either the 5′ carbon of 5′-dA, or the C5 carbon of tRNA U34. **(B)** Comparison of WT Elp3 activity with AcCoA to E207, K302, and Y304 mutants and a -SAM negative control reveals that E207A, K302R, K302A, and Y304A display no or nearly no activity with AcCoA, while Y304F retains partial activity. **(C)** Comparison of WT Elp3 activity with acetate to E207, K302, and Y304 mutants and a -SAM negative control reveals that all of these mutants are inactive with acetate. Activity assays in **(B)** and **(C)** were performed in triplicate with 5 µM Elp3, 4.4 µM tRNA Arg^UCU^, 0.5 mM dithionite as reductant, and either 27.5 µM AcCoA or 2 mM acetate, with endpoints collected after 4 hrs. Errors are shown as SD.

To test these predictions, we mutated the conserved rSAM active site residues E207, K302, and Y304 (**Figure S12**), which make hydrogen bond contacts with acetate in the docking models, and measured their relative tRNA modifying activities. In reactions with ^14^C-AcCoA (**Figure 4B**), all mutations to E207 (E207A) or K302 (K302R, K302A) resulted in nearly complete loss of tRNA modifying activity relative to WT Elp3. For Y304, while the Ala mutant Y304A resulted in complete loss of activity, the Phe mutant Y304F retained about 30% activity relative to WT, showing that while the Y304 hydroxyl group is important for Elp3 reactivity with AcCoA, it is not essential. We next similarly tested reactions with ^14^C-acetate and the K302(R or A) and Y304(F or A) mutations (**Figure 4C**). In the case of acetate-based Elp3 reactivity, all of the tested mutants were inactive. These mutational data show that both AcCoA- and acetate-based Elp3 reactivity are critically sensitive to the same rSAM active site mutations, consistent with our proposed mechanism in which molecular acetate is a key reaction intermediate in the Elp3-catalyzed tRNA modification reaction. The lower activity observed for acetate- versus AcCoA-based reactivity with Elp3 mutant Y304F may reflect the fact that reactions carried out with AcCoA, where acetate is released into the enclosed molecular tunnel, are likely able to achieve a dramatically higher local concentration of acetate in the rSAM active site, compared to reactions carried out with only exogenous acetate. Indeed, with a calculated Elp3 tunnel volume of 370 ± 30 Å^3^, one molecule of acetate released into the enclosed molecular tunnel would have an effective concentration of approximately 4.5 M; this is much higher than the 2 mM ^14^C-acetate added during reactions with only exogenous acetate, likely making acetate-based reactivity more sensitive to mutations that weaken its binding in the rSAM active site.

## Discussion

Elp3 and Elongator install the key cm^5^U34 modification on the anticodon loop of tRNA, a critical intermediate for the subsequent formation of a whole family of cm^5^U-derived tRNA modifications (**Figure S1**) that have diverse impacts on cellular translational efficiency and fidelity across all domains of life. Over the past decade, multiple mechanisms have been proposed to explain how this functionally important tRNA modification is installed by the rSAM and KAT domains of Elp3. In 2014, Selvadurai *et al* proposed two possible, general mechanisms for Elp3-mediated tRNA modification based on isotope incorporation experiments (**Figure 5**): (A) 5′-dA· generated at the rSAM active site abstracts a hydrogen atom directly from AcCoA to form an AcCoA-based radical that subsequently reacts with tRNA U34 resulting in an AcCoA-tRNA covalent intermediate that is hydrolyzed to ultimately produce cm^5^U (**Figure 5A**). Alternatively, (B) AcCoA first acetylates an unspecified Elp3 residue, 5′-dA· abstracts a hydrogen atom from the acetylated Elp3 residue to generate an acetyl-based radical, which reacts with tRNA U34 to produce a covalent Elp3-tRNA adduct, and finally hydrolysis of the acetylated Elp3 residue produces cm^5^U (**Figure 5B**). Since 2019, multiple structures of Elp3 and Elongator subunits in complex with tRNA have been determined,^30,34,35^ and based on this structural data Abbassi *et al.* have more recently proposed a variation on mechanism (B) above, in which (C): AcCoA acetylates a conserved Elp3 lysine residue, the acetyl group is somehow passed from residue to residue along Elp3, to ultimately produce an acetylated lysine (*Min* K302, human K316; **Table S1**) in the rSAM active site of Elp3. In this mechanism, 5′-dA· is proposed to abstract a hydrogen atom from an active site tyrosine (*Min* Y304, human Y318; **Table S1**), the resulting tyrosyl radical reacts with the acetylated active site lysine, and the acetyl lysine radical then reacts with tRNA U34 (**Figure 5C**).^35^ However, in large part due to challenges in obtaining active recombinant Elp3, none of these mechanisms have been subjected to enzymological investigations *in vitro* that might more clearly define the biochemical mechanism of Elp3- and Elongator-mediated tRNA modification.

**Figure 5.**
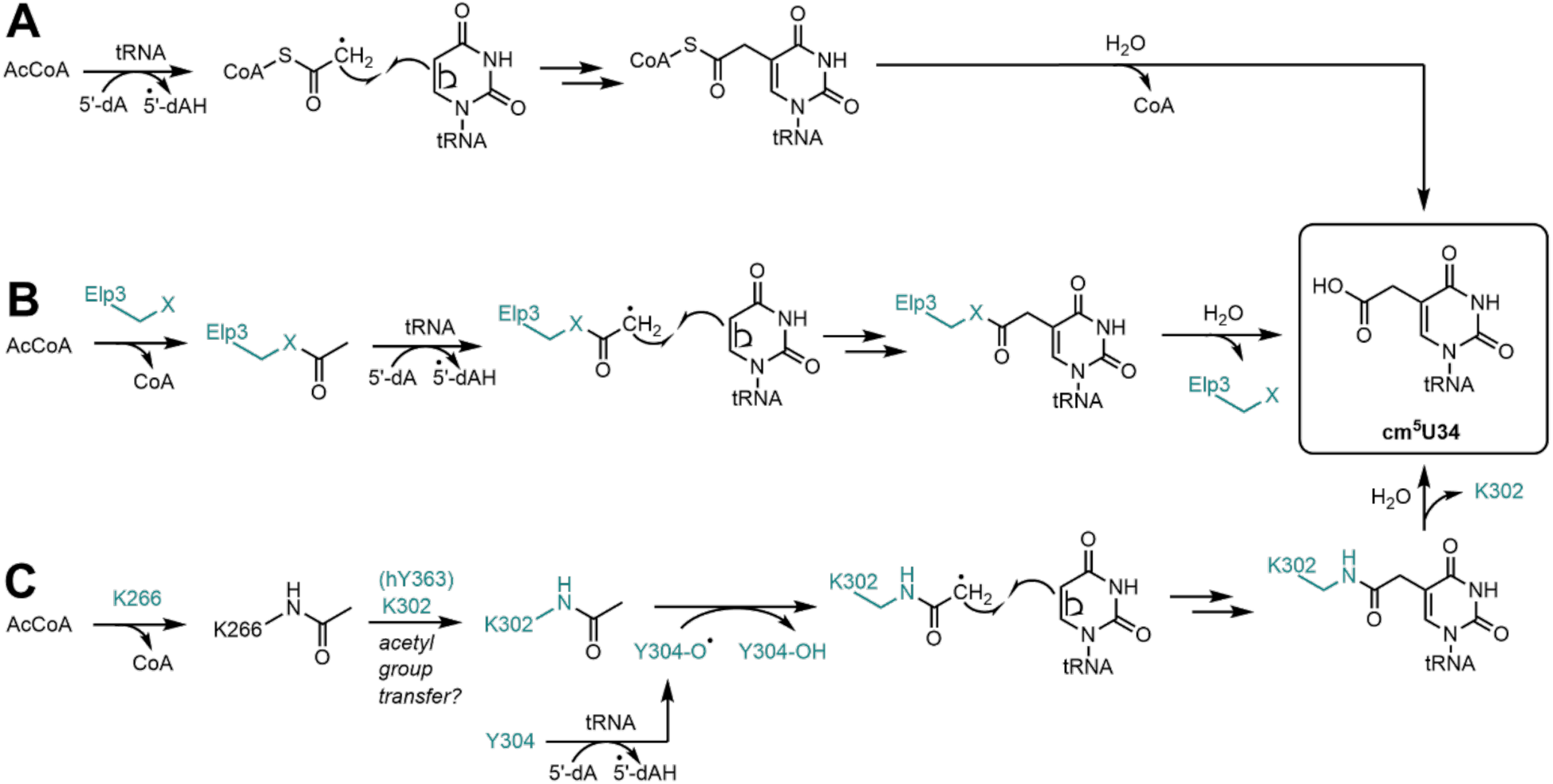
Previously proposed Elp3 tRNA modification mechanisms. **(A)** and **(B)** are mechanisms originally proposed by Selvadurai *et al* in 2014, characterized by either reaction of AcCoA directly with 5′-dA· **(A)**, or initial acetylation of Elp3 by AcCoA followed by reaction of the acetylated Elp3 residue with 5′-dA· **(B)**. **(C)** is a variation on (B) more recently proposed by Abassi *et al* in 2024, in which AcCoA is used to acetylate a conserved lysine, this covalent acetyl group is then transferred from residue to residue along Elp3, to ultimately react with a tyrosyl radical in the rSAM active site.

Here we have chemically reconstituted the [4Fe-4S] cluster and obtained robust *in vitro* tRNA modification activity for a model, recombinant archaeal Elp3. Our structural analysis of Elp3-tRNA complexes revealed a conserved molecular tunnel connecting the rSAM and KAT active sites on Elp3 (**Figure 1A**), suggesting a new possible mechanism for Elp3-mediated tRNA modification where the molecular tunnel might be used to transport free acetate (or acetic acid) molecules between the distant Elp3 active sites. In support of this hypothesis, our *in vitro* kinetic assays with reconstituted Elp3 showed that acetate can be used in place of AcCoA to carry out cm^5^U formation on tRNA, and that the CoA portion of AcCoA is not needed to carry out the tRNA modification reaction at sufficient acetate concentrations (**Figure 1 B-D**). Sterically blocking the molecular tunnel by mutagenesis disrupts Elp3-mediated tRNA modification activity without significantly affecting tRNA binding or AcCoA binding and turnover (**Figure 2**). Similar experiments mutating tunnel-lining residues to either block or open tunnels have been used to verify and elucidate the functions of molecular tunnels that transport small molecules or gases in variety of other enzymes.^53–59^ Based on our structural analysis and biochemical data, we therefore propose a new model (**Figure 6**) for tRNA modification by Elp3 and Elongator whereby the acetyl group from AcCoA is delivered to the rSAM active site via transport of free acetate through a conserved, 20+ Å-long molecular tunnel connecting both domains of Elp3.

**Figure 6.**
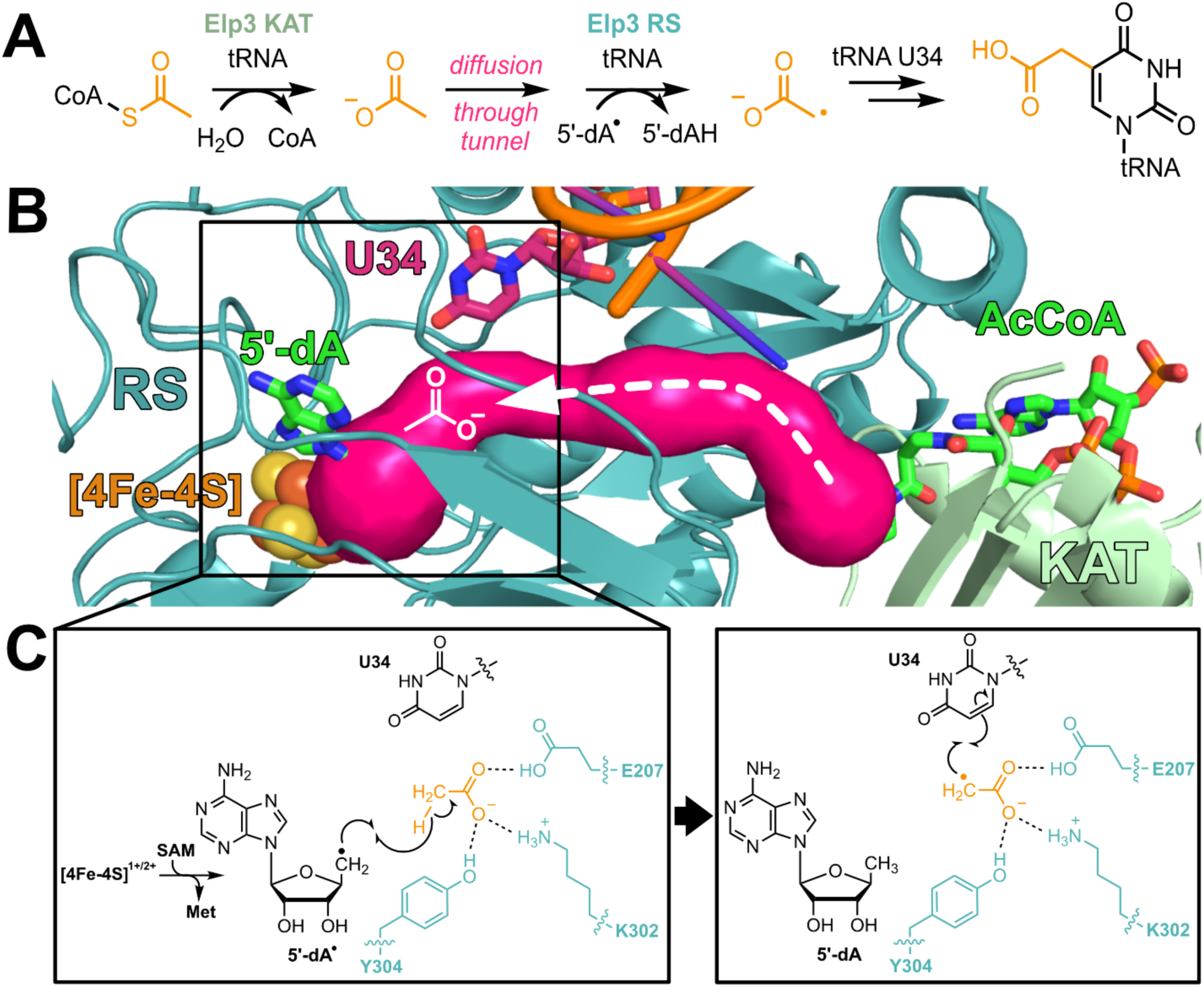
Summary of Elp3 tRNA modification reaction mechanism involving transport of acetate through the Elp3 molecular tunnel. **(A)** Overview of proposed Elp3 tRNA modification reaction. Elp3 hydrolyzes AcCoA in the KAT site to produce acetate (or acetic acid, depending on local pH), which then diffuses through Elp3’s molecular tunnel to the rSAM active site. Once acetate is in the rSAM site, 5′-dA· abstracts a hydrogen from acetate to produce an acetate radical, which adds to the C5 position of U34 on substrate tRNA to form cm^5^U. **(B)** Structural overview of proposed Elp3 tRNA-modifying reaction using the yeast structure of Elp3 bound to tRNA (PDB 8ASW). The Elp3 rSAM domain is colored teal, the Elp3 KAT domain is colored light green, and the calculated tunnel is shown in pink; 5′-dA and AcCoA analog desulfo-CoA (aligned from PDB 6IA6) are shown in green sticks. tRNA and substrate tRNA base U34 are shown in orange and pink, respectively. **(C)** Schematic of acetate’s reaction with 5′-dA· and U34 in the rSAM active site. Left: key conserved rSAM active site residues (teal) noncovalently position acetate (orange) for hydrogen atom abstraction by 5′-dA·. Right: next, the coordinated acetate radical adds to the C5 position of substrate tRNA U34 to ultimately form cm^5^U-modified tRNA. All Elp3 residues are shown with *Min* numbering; see **Table S1** for eukaryotic residue numbering.

Approximately a dozen known enzymes use a molecular tunnel to connect two distant active sites within the enzyme, ^58,60–62^ most commonly for the transport ammonia,^61^ though examples of indole,^63^ carbon monoxide,^64^ dehydroglycine,^62,65^ and acetaldehyde^66^ transport are known. Multisite tunnel enzymes appear to have evolved their tunnels independently from one another, though some share convergent similarities, such as a similar makeup of tunnel-lining residues and high conservation of these residues, as well as similar tunnel lengths of ∼20-30 Å.^61^ To our knowledge, Elp3 and Elongator would be the only known enzyme or enzymatic complex to use a tunnel for the transport of molecular acetate between multiple active sites. The Elp3 tunnel is composed largely from its partial TIM barrel motif, a feature it shares in common with imidazole glycerolphosphate synthase (IGPS)^67,68^ and fellow radical SAM enzyme HydG,^62,65^ both of which use full TIM barrels to transport intermediates, while Elp3’s partial TIM barrel is capped by tRNA to complete the tunnel. That multiple enzymes utilize a TIM barrel or partial TIM barrel in their molecular tunnels is hardly surprising, as the tubular interior of the motif’s architecture readily lends itself to small molecule or ion transport. Across the huge radical SAM enzyme superfamily however, Elp3 would be only the third identified rSAM enzyme that uses a molecular tunnel to transport substrates or intermediates; the only other known examples being HydG, mentioned above, and the closely related enzyme ThiH, both of which transport dehydroglycine as a reaction intermediate using enclosed tunnels.^62^ Elp3 is the only example where the tunnel is formed in part by RNA, suggesting that other rSAM enzymes may form complexes that generate molecular tunnels. The rSAM superfamily is incredibly diverse, with over 700,000 unique sequences identified,^69^ less than 4000 of which have been functionally characterized^70^ and less than 100 of which have been structurally characterized.^71,72^ We therefore anticipate that other tunnel-using rSAM enzymes likely exist and have yet to be discovered due the large, but largely unexplored, diversity of radical SAM enzymes and their conserved architecture featuring a full or partial TIM barrel motif.^72,73^

Molecular docking simulations using CaverDock and SeamDock to model possible acetate positions within the Elp3 molecular tunnel suggested acetate poses in the Elp3 rSAM active site where the acetate methyl group is positioned directly between the 5′ carbon of 5′-dA and the C5 carbon of tRNA U34, at distances reasonable for radical-mediated reactivity between each of these reaction components (**Figure 4A, Figure S11**). This docking pose also places acetate within hydrogen bonding distance to key conserved, active site residues E207, K302, and Y304. Mutating these rSAM active site residues results in inactivation of Elp3 for AcCoA- and acetate-based reactions in all cases except for the Y304F mutation with AcCoA, which retains partial activity (**Figures 4B & 4C**). These data are consistent with previous mutagenesis experiments in yeast-based assays, showing K302 and Y304 are important for Elongator function,^30,34,35^ additionally reveal the importance of E207, and show that AcCoA- and acetate-based reactivity are similarly dependent on the same subset of rSAM active site residues. Our observation that the Y304F mutation retains partial activity with AcCoA is also consistent with previous isotope incorporation experiments showing direct reaction of the AcCoA acetyl group with 5′-dA·,^31^ in contrast to recent proposals suggesting intermediate radical transfer through Y304 (human Y318), which would strictly require the tyrosyl hydroxyl group.^30,35^ Finally, our isotope incorporation experiments showing that reaction of doubly ^18^O-labeled acetate results in formation of doubly ^18^O-labeled cm^5^U suggest that acetate is not required to undergo nucleophilic or hydrolytic reaction during Elp3’s catalytic cycle, further supporting the idea that Elp3 transports and uses free acetate rather than an acetylated residue intermediate.

Together, our biochemical data and structural analyses show how Elp3 and Elongator use a conserved molecular tunnel to transport acetate from the Elp3 KAT domain to the rSAM active site, where acetate is likely positioned through key noncovalent interactions to undergo reaction with 5′-dA· and then tRNA U34 to catalyze formation of the cm^5^U modification on tRNA. Our proposed mechanism for Elp3- and Elongator-mediated tRNA modification answers long-standing questions about how the enzyme bridges the large distance between the AcCoA binding site on the KAT domain and the [4Fe-4S]-containing active site in the rSAM domain. This provides important groundwork toward understanding how tRNA and other Elp proteins in the Elongator complex may help regulate tRNA modification chemistry and how the active site of Elp3 could be targeted with small molecules mimicking acetate. More broadly, this work elucidates a novel tRNA modification and acetate transport mechanism and suggests that the rSAM enzyme superfamily may use diverse, yet-to-be-discovered molecular tunnels to carry out intermediate transport and reactivity.

## Methods

### CAVER, CaverDock, and SeamDock analysis

Analysis of the tRNA-bound yeast Elp3 structure (from PDB 8ASW)^30^ was performed with the CAVER 3.0.3 Pymol plugin.^48^ CAVER settings were as follows: probe radius 1.4 Å, starting coordinates x=186.918, y=158.706, z=197.689, and input atoms included all 20 amino acids and all 4 RNA bases, but not 5′-dA or the [4Fe-4S] cluster. The model of yeast Elp3 with tRNA bound derived from PDB 8ASW and the CAVER tunnel generated using the above were used as CaverDock inputs. A model of acetate was obtained from the PDB.^71^ The CAVER tunnel was discretized into discs with 0.3 Å between individual discs and the direction of tunnel analysis was reversed to model binding the acetate ligand from the tunnel’s entrance at the KAT site into the interior of Elp3. SeamDock analysis was performed on of the tRNA-bound yeast Elp3 structure and an acetate model obtained from the PDB. The box center coordinates were set to x=7, y=-6, z=3 and all other settings were set to defaults.

### In vitro transcription (IVT) of Elp3 tRNA substrate

The *Methanocaldococcus infernus* (*Min*) tRNA^Arg^UCU sequence (5′- GGACCCGUAGCCUAGCCUGGAUAGGGCACCGGCCUUCUAAGCCGGGGGUCGGGGGUUCAAAUCCCCCCGGGUCCGCCA-3′) was obtained from sequences deposited in the Genomic tRNA Database.^74^ The corresponding tRNA^Arg^UCU DNA template was PCR-amplified from a commercially-obtained DNA oligonucleotide reverse complement sequence containing a 5′ T7 promoter sequence (Integrated DNA Technologies; template DNA sequence = 5′- TGGCGGACCCGGGGGGATTTGAACCCCCGACCCCCGGCTTAGAAGGCCGGTGCCCTATCCAGGCTAGGCTACGGGTCCTATAGTGAGTCGTATTA-3′). The tRNA was *in vitro* transcribed using T7 polymerase in 500 µL reactions: 0.25 µM DNA template, 2.5 mM rNTPs, 0.05 % Triton X-100 (v/v), 5 µL of 10 mg/mL T7 polymerase, and 1 Unit of thermostable inorganic pyrophosphatase (New England Biolabs) in T7 buffer (40 mM Tris HCl, pH 7.5, 50 mM MgCl2, 2 mM spermidine, 5 mM dithiothreitol (DTT)). Reactions were incubated at 37 °C for 3 hr, then the DNA template was digested with TURBO DNase (Thermo Fisher Scientific) for 1 hr at 37 °C. Following this, tRNA was purified using denaturing urea PAGE gels, phenol:chloroform extracted, and precipitated with 80% ethanol at -20 °C. Isolated tRNA was redissolved in water, annealed for 2 min at 80 °C and 2 min at 60 °C, then MgCl2 was added to a final concentration of 10 mM and the tRNA was aliquoted, flash frozen with liquid nitrogen, and stored at -70 °C until ready for use.

### Cloning and overexpression of recombinant Min Elp3

The *Min* Elp3 gene was ordered as an *E. coli*-codon optimized DNA sequence (Integrated DNA Technologies) and cloned into a pET28a-tev vector containing an N-terminal 6xHis affinity tag and tobacco etch virus protease (TEV) cleavage site using Gibson assembly (New England Biolabs HiFi DNA Assembly Kit). The final expression vector sequence was confirmed by whole plasmid DNA sequencing (Plasmidsaurus). Elp3 point mutations were generated using site-directed mutagenesis by whole-plasmid polymerase chain reaction (PCR) and confirmed by DNA sequencing (Plasmidsaurus).

*Min* Elp3 plasmids and pDB1282 plasmid containing the *isc* operon were simultaneously transformed into *E. coli* BL21(DE3) cells. Bacterial growth was carried out in LB media at 37 °C with shaking. At an optical density of 600 nm (OD600) of 0.3-0.4, the *isc* operon was induced with a 20 % (w/v) solution of L-(+)-arabinose for a final concentration of 0.2 % (w/v). Solutions of 200 mM iron(III) chloride and 200 mM cysteine were added to a final concentration of 0.2 mM each. At an OD600 of 0.6-0.8, Elp3 expression was induced by adding 1 M IPTG to a final concentration of 1 mM. Induction occurred at 18 °C overnight with shaking. Cells were harvested at 6000 x *g* for 30 min at 4 °C, flash frozen with liquid nitrogen, and stored at -70 °C until ready for use.

### Purification of Min Elp3

Purification of *Min* Elp3 constructs was mostly carried out in an anaerobic chamber (MBraun) at an oxygen concentration of <1 ppm and temperature of 16 °C. Buffers used during purification were as follows: lysis/wash buffer (50 mM HEPES, pH 7.5, 300 mM KCl, 2 mM imidazole, 10 % glycerol (v/v)), wash 2 buffer (50 mM HEPES, pH 7.5, 1.0 M KCl, 10 mM imidazole, 10 % glycerol (v/v)), wash 3 buffer (50 mM HEPES, pH 7.5, 300 mM KCl, 10 mM imidazole, 10 % glycerol (v/v)), elution buffer (50 mM HEPES, pH 7.5, 300 mM KCl, 250 mM imidazole, 10 % glycerol (v/v)), size exclusion chromatography (SEC) buffer (50 mM HEPES, pH 7.5, 300 mM KCl). All buffers were thoroughly deoxygenated by bubbling with argon for 1 hr before being introduced into the anaerobic chamber and then stirred vigorously overnight to equilibrate. 10 mM 2-mercaptoethanol (BME) was added to buffers before use, with the exception of SEC buffer, which had 5 mM DTT added. Cell pellets were thawed inside the anaerobic chamber and resuspended in 35 mL of lysis/wash buffer, then lysed by sonication on ice using an air-tight sonicator adaptor outside of the anaerobic chamber (Branson Sonifier 450, 4 x 2 min cycles at setting 6, 50 % duty cycle). Following sonication, lysate was brought back into the anaerobic chamber and transferred to centrifuge tubes which were sealed and centrifuged outside of the anaerobic chamber for 45 min at 14,500 x *g* and 4 °C. Sealed centrifuge tubes were brought back into the anaerobic chamber and lysate was mixed for 1 hr with Ni-NTA resin that had been deoxygenated by bubbling with argon gas and equilibrated with lysis/wash buffer. The mixture was transferred to a gravity column and the flow-through was collected. The resin was then washed with 50 mL of wash 2 buffer and 50 mL of wash 3 buffer, which were collected, before protein was eluted with 15 mL of elution buffer in 1 mL fractions. Following identification of Elp3-containing fractions by gel electrophoresis (10 % polyacrylamide gel), fractions were pooled and concentrated to ∼2 mL using Amicon Ultra 0.5 mL spin columns (50 kDa cutoff) and subjected to size exclusion chromatography on an AKTA Pure 25M with a 16/60 S-200pg column, pre-equilibrated with 3 column volumes of deoxygenated sizing buffer. SEC fractions were collected inside the anaerobic chamber.

### Chemical reconstitution of Min Elp3

Following identification of desired SEC fractions by gel electrophoresis (10 % polyacrylamide gel), chosen fractions were concentrated and exchanged into reconstitution buffer (100 mM HEPES, pH 7.5, 500 mM KCl, 10 % glycerol) using Amicon Ultra 0.5 mL spin columns (50 kDa cutoff) inside the anaerobic chamber. The purified Elp3 was then diluted to 75 µM in reconstitution buffer, with DTT added to a final concentration of 5 mM, and incubated for 1 hr. Iron(III) chloride was gradually added to a 10-fold molar excess relative to Elp3 (6 aliquots, with 5 min between aliquot additions) and after 10 min of further incubation, sulfide (Na2S) was gradually added to a 10-fold molar excess (6 aliquots, with 15 min between aliquot additions). The reconstitution was allowed to proceed overnight. All reconstitution steps were carried out at 4 °C in the anaerobic chamber. Following reconstitution, Elp3 was centrifuged to remove any precipitate (25 min at 14,500 rpm, Eppendorf MiniSpin plus), buffer exchanged into reaction buffer (50 mM HEPES, pH 7.5, 150 mM KCl, 300 mM NaCl, 5 mM MgCl2, 1 % glycerol), concentrated, glycerol added to a final concentration of 20 %, and flash frozen with liquid nitrogen. Elp3 aliquots were stored at -70 °C for further use. UV-Vis spectra were collected to verify the presence of the characteristic [4Fe-4S] absorption peak at 420 nm (Thermo Scientific NanoDrop 2000c).

### In vitro Elp3 activity assays

Substrate tRNA aliquots were brought into the anaerobic chamber and deoxygenated by equilibrating with the anaerobic atmosphere at 4 °C for 30 min. A 2X concentration Elp3 solution was prepared at 10 µM with 55 µM AcCoA, 50 µM SAM, 1 mM sodium dithionite, and 2X reaction buffer and incubated for 10 min at 16 °C. A 2X concentration tRNA solution was prepared by diluting tRNA aliquots to 8.8 µM with water. Reactions were initiated by mixing the 2X Elp3 and 2X tRNA solutions and incubated at 37 °C. Final reaction concentrations were 5 µM Elp3, 27.5 µM AcCoA, 25 µM SAM, 0.5 mM dithionite, and 4.4 µM tRNA. Activity assays with acetate were carried out similarly, where the relevant concentration of acetate was used in place of AcCoA. 10 µL timepoint aliquots were quenched with 1 µL of 10 mM ethylenediaminetetraacetic acid (EDTA) stock solution. For each quenched timepoint, tRNA was then isolated from the other reaction components (Zymo RNA Clean and Concentrator kit) and eluted in 15 µL water. For radiolabel-based assays, 12 µL eluted tRNA was added to 10 mL of scintillation fluid (Ultima Gold) in a 20 mL glass scintillation vial and counted on a liquid scintillation counter (PerkinElmer Quantulus GCT 6220). For mass spectrometry-based assays, eluted tRNA was digested into individual nucleosides (NEB Nucleoside Digestion Mix) and analyzed by LC-MS or LC-MS/MS. For the ^18^O isotope incorporation experiment, the activity assay was performed with 25 µM Elp3, 10 mM ^18^O2- or normal abundance acetate, 50 µM SAM, 0.5 mM dithionite, and 4.4 µM tRNA.

### Electrophoretic mobility shift assays (EMSAs)

EMSAs were based on a previously published procedure for Elp3.^75^ 250 nM tRNA was incubated with a 1:2 serial dilution of 20 µM to 39 nM Elp3, plus 0 µM Elp3 control, in incubation buffer (100 mM HEPES, pH 7.5, 750 mM NaCl, 25 mM DTT) at 20 °C for 30 min. Non-denaturing 5 % TBE gels (BioRad) were pre-run in running buffer (3.3 g Tris, 14.4 g glycine, 150 mg DTT per L) at 5 mA for 1 hr at 4 °C. 10 µL of the incubated Elp3- tRNA samples were mixed with 2 µL of 50 % glycerol in incubation buffer and loaded onto the gel. Gels were run at 5 mA for ∼2 hr with fresh running buffer, stained with Sybr Gold (Thermo Fisher Scientific) for 30 min, and imaged (Cell Biosciences FluorChem Q). Gel images were analyzed using ImageJ.^76^

### Acetyl-CoA hydrolysis assays

In an anaerobic chamber, triplicate 2X concentration solutions of Elp3 and AcCoA were incubated at 16 °C for 10 min in 2X reaction buffer, then mixed with a 2X concentration solution of tRNA to initiate AcCoA hydrolysis at a final reaction concentration of 10 µM Elp3, 100 µM AcCoA, and 2 µM tRNA. Reactions were then incubated at 37 °C for 20 min and quenched by freezing. AcCoA consumed in each reaction tube was determined using a fluorescence-based AcCoA assay kit (Sigma MAK039) and measured on a TECAN Spark plate reader.

## Acknowledgements

We would like to thank Kaitlyn Tsai and Danica Fujimori at UCSF for gifting us the isc operon plasmid and initial advice on rSAM reconstitution protocols. This material is based upon work supported by the National Science Foundation under Grant No. 2339759. This work was also funded by the University of Delaware Research Foundation.

## References

1. Schultz, S. K.; Kothe, U. RNA modifying enzymes shape tRNA biogenesis and function. Journal of Biological Chemistry 2024, 300, 107488.

2. Suzuki, T. The expanding world of tRNA modifications and their disease relevance. Nature Reviews Molecular Cell Biology 2021, 22, 375–392.

3. Yared, M.; Marcelot, A.; Barraud, P. Beyond the Anticodon: tRNA Core Modifications and Their Impact on Structure, Translation and Stress Adaptation. Genes 2024, 15, 374.

4. Smith, T. J.; Giles, R. N.; Koutmou, K. S. Anticodon stem-loop tRNA modifications influence codon decoding and frame maintenance during translation. Seminars in Cell & Developmental Biology 2023, *154B*, 105–113.

5. Cappannini, A.; Ray, A.; Purta, E.; Mukherjee, S.; Boccaletto, P.; Moafinejad, S. N.; Lechner, A.; Barchet, C.; Klaholz, B. P.; Stefaniak, F.; Bujnicki, J. M. MODOMICS: a database of RNA modifications and related information. 2023 update. Nucleic Acids Research 2023, 52, D239–D244.

6. Pan, T. Modifications and functional genomics of human transfer RNA. Cell Research 2018, 28, 395–404.

7. Machnicka, M. A.; Olchowik, A.; Grosjean, H.; Bujnicki, J. M. Distribution and frequencies of post-transcriptional modifications in tRNAs. RNA Biology 2015, 11, 1619–1629.

8. Schaffrath, R.; Leidel, S. A. Wobble uridine modifications–a reason to live, a reason to die?! RNA Biology 2017, 14, 1209–1222.

9. Rezgui, V. A. N.; Tyagi, K.; Ranjan, N.; Konevega, A. L.; Mittelstaet, J.; Rodnina, M. V.; Peter, M.; Pedrioli, P. G. A. tRNA tK UUU, tQ UUG, and tE UUC wobble position modifications fine-tune protein translation by promoting ribosome A-site binding. Proc. Natl. Acad. Sci. U. S. A. 2013, 110, 12289–12294.

10. Vendeix, F. A. P.; Murphy, F. V.; Cantara, W. A.; Leszczyńska, G.; Gustilo, E. M.; Sproat, B.; Malkiewicz, A.; Agris, P. F. Human tRNALys3UUU Is Pre-Structured by Natural Modifications for Cognate and Wobble Codon Binding through Keto–Enol Tautomerism. J. Mol. Biol. 2012, 416, 467–485.

11. Ramos, J.; Fu, D. The emerging impact of tRNA modifications in the brain and nervous system. Biochimica et Biophysica Acta - Gene Regulatory Mechanisms 2019, 1862, 412–428.

12. Begley, U.; Dyavaiah, M.; Patil, A.; Conklin, D. S.; Zitomer, R. S.; Begley, T. J. Trm9- Catalyzed tRNA Modifications Link Translation to the DNA Damage Response. Molecular Cell 2007, 28, 860–870.

13. Bennett, C. B.; Lewis, L. K.; Karthikeyan, G.; Lobachev, K. S.; Jin, Y. H.; Sterling, J. F.; Snipe, J. R.; Resnick, M. A. Genes required for ionizing radiation resistance in yeast. Nat Genet 2001, 29, 426–434.

14. Patil, A.; Dyavaiah, M.; Joseph, F.; Rooney, J. P.; Chan, C. T. Y.; Dedon, P. C.; Begley, T. J. Increased tRNA modification and gene-specific codon usage regulate cell cycle progression during the DNA damage response. Cell Cycle 2012, 11, 3656–3665.

15. Fu, D.; Brophy, J. A. N.; Chan, C. T. Y.; Atmore, K. A.; Begley, U.; Paules, R. S.; Dedon, P. C.; Begley, T. J.; Samson, L. D. Human AlkB Homolog ABH8 Is a tRNA Methyltransferase Required for Wobble Uridine Modification and DNA Damage Survival. Molecular and Cellular Biology 2010, 30, 2449–2459.

16. Endres, L.; Begley, U.; Clark, R.; Gu, C.; Dziergowska, A.; Małkiewicz, A.; Melendez, J. A.; Dedon, P. C.; Begley, T. J. Alkbh8 Regulates Selenocysteine-Protein Expression to Protect against Reactive Oxygen Species Damage. PLoS ONE 2015, 10, e0131335.

17. Lee, M. Y.; Ojeda-Britez, S.; Ehrbar, D.; Samwer, A.; Begley, T. J.; Melendez, J. A. Selenoproteins and the senescence-associated epitranscriptome. Experimental Biology and Medicine 2022, 247, 2090–2102.

18. Tükenmez, H.; Xu, H.; Esberg, A.; Byström, A. S. The role of wobble uridine modifications in +1 translational frameshifting in eukaryotes. Nucleic Acids Res 2015, 43, 9489–9499.

19. Karlsborn, T.; Mahmud, A K M Firoj; Tükenmez, H.; Byström, A. S. Loss of ncm5 and mcm5 wobble uridine side chains results in an altered metabolic profile. Metabolomics 2016, 12, 177.

20. Deng, W.; Babu, R.; Su, D.; Yin, S.; Begley, T. J.; Dedon, P. C. Trm9-Catalyzed tRNA Modifications Regulate Global Protein Expression by Codon-Biased Translation. PLOS Genetics 2015, 11, e1005706.

21. Klassen, R.; Grunewald, P.; Thüring, K. L.; Eichler, C.; Helm, M.; Schaffrath, R. Loss of Anticodon Wobble Uridine Modifications Affects tRNALys Function and Protein Levels in Saccharomyces cerevisiae. PLOS One 2015, 10, e0119261.

22. Klassen, R.; Ciftci, A.; Funk, J.; Bruch, A.; Butter, F.; Schaffrath, R. tRNA anticodon loop modifications ensure protein homeostasis and cell morphogenesis in yeast. Nucleic Acids Res 2016, 44, 10946–10959.

23. Karlsborn, T.; Tükenmez, H.; Chen, C.; Byström, A. S. Familial dysautonomia (FD) patients have reduced levels of the modified wobble nucleoside mcm5s2U in tRNA. Biochemical and Biophysical Research Communications 2014, 454, 441–445.

24. Huang, B.; Johansson, M. J. O.; Byström, A. S. An early step in wobble uridine tRNA modification requires the Elongator complex. RNA 2005, 11, 424–436.

25. Songe-Møller, L.; van den Born, E.; Leihne, V.; Vågbø, C. B.; Kristoffersen, T.; Krokan, H. E.; Kirpekar, F.; Falnes, P. Ø; Klungland, A. Mammalian ALKBH8 Possesses tRNA Methyltransferase Activity Required for the Biogenesis of Multiple Wobble Uridine Modifications Implicated in Translational Decoding. Molecular and Cellular Biology 2010, 30, 1814–1827.

26. van den Born, E.; Vågbø, C. B.; Songe-Møller, L.; Leihne, V.; Lien, G. F.; Leszczynska, G.; Małkiewicz, A.; Krokan, H. E.; Kirpekar, F.; Klungland, A.; Falnes, P. Ø ALKBH8-mediated formation of a novel diastereomeric pair of wobble nucleosides in mammalian tRNA. Nature Communications 2011, 2, 172.

27. Nakai, Y.; Nakai, M.; Yano, T. Sulfur Modifications of the Wobble U34 in tRNAs and their Intracellular Localization in Eukaryotic Cells. Biomolecules 2017, 7, 17.

28. Kogaki, T.; Hase, H.; Tanimoto, M.; Tashiro, A.; Kitae, K.; Ueda, Y.; Jingushi, K.; Tsujikawa, K. ALKBH4 is a novel enzyme that promotes translation through modified uridine regulation. J. Biol. Chem. 2023, 299, 105093.

29. Dauden, M. I.; Kosinski, J.; Kolaj-Robin, O.; Desfosses, A.; Ori, A.; Faux, C.; Hoffmann, N. A.; Onuma, O. F.; Breunig, K. D.; Beck, M.; Sachse, C.; Séraphin, B.; Glatt, S.; Müller, C. W. Architecture of the yeast Elongator complex. EMBO Reports 2017, 18, 264–279.

30. Jaciuk, M.; Scherf, D.; Kaszuba, K.; Gaik, M.; Rau, A.; Kościelniak, A.; Krutyhołowa, R.; Rawski, M.; Indyka, P.; Graziadei, A.; Chramiec-Głąbik, A.; Biela, A.; Dobosz, D.; Lin, T.; Abbassi, N.; Hammermeister, A.; Rappsilber, J.; Kosinski, J.; Schaffrath, R.; Glatt, S. Cryo-EM structure of the fully assembled Elongator complex. Nucleic Acids Research 2023, gkac1232.

31. Selvadurai, K.; Wang, P.; Seimetz, J.; Huang, R. H. Archaeal Elp3 catalyzes tRNA wobble uridine modification at C5 via a radical mechanism. Nature Chemical Biology 2014, 10, 810–812.

32. Glatt, S.; Zabel, R.; Kolaj-Robin, O.; Onuma, O. F.; Baudin, F.; Graziadei, A.; Taverniti, V.; Lin, T.; Baymann, F.; Séraphin, B.; Breunig, K. D.; Müller, C. W. Structural basis for tRNA modification by Elp3 from Dehalococcoides mccartyi. Nature Structural & Molecular Biology 2016, 23, 794–802.

33. Lin, T.; Abbassi, N. E. H.; Zakrzewski, K.; Chramiec-Głąbik, A.; Jemioła-Rzemińska, M.; Różycki, J.; Glatt, S. The Elongator subunit Elp3 is a non-canonical tRNA acetyltransferase. Nature Communications 2019, 10, 625.

34. Dauden, M. I.; Jaciuk, M.; Weis, F.; Lin, T.; Kleindienst, C.; Abbassi, N. E. H.; Khatter, H.; Krutyhołowa, R.; Breunig, K. D.; Kosinski, J.; Müller, C. W.; Glatt, S. Molecular basis of tRNA recognition by the Elongator complex. Science Advances 2019, 5, eaaw2326.

35. Abbassi, N.; Jaciuk, M.; Schert, D.; Böhnert, P.; Rau, A.; Hammermeister, A.; Rawski, M.; Indyka, P.; Wazny, G.; Chramiec-Głąbik, A.; Dobosz, D.; Skupien-Rabian, B.; Janowska, U.; Rappsilber, J.; Schaffrath, R.; Lin, T.; Glatt, S. Cryo-EM structures of the human Elongator complex at work. 2024, 15, 4094.

36. Pereira, M.; Francisco, S.; Varanda, A. S.; Santos, M.; Santos, M. A. S.; Soares, A. R. Impact of tRNA Modifications and tRNA-Modifying Enzymes on Proteostasis and Human Disease. International Journal of Molecular Sciences 2018, 19, 3738.

37. Nedialkova, D. D.; Leidel, S. A. Optimization of Codon Translation Rates via tRNA Modifications Maintains Proteome Integrity. Cell 2015, 161, 1606–1618.

38. Ladang, A.; Rapino, F.; Heukamp, L. C.; Tharun, L.; Shostak, K.; Hermand, D.; Delaunay, S.; Klevernic, I.; Jiang, Z.; Jacques, N.; Jamart, D.; Migeot, V.; Florin, A.; Göktuna, S.; Malgrange, B.; Sansom, O. J.; Nguyen, L.; Büttner, R.; Close, P.; Chariot, A. Elp3 drives Wnt-dependent tumor initiation and regeneration in the intestine. Journal of Experimental Medicine 2015, 212, 2057–2075.

39. Wang, Y.; Tao, E.; Tan, J.; Gao, Q.; Chen, Y.; Faing, J. tRNA modifications: insights into their role in human cancers. Trends in Cell Biology 2023, 33, 1035–1048.

40. Rapino, F.; Delaunay, S.; Rambow, F.; Zhou, Z.; Tharun, L.; De Tullio, P.; Sin, O.; Shostak, K.; Schmitz, S.; Piepers, J.; Ghesquière, B.; Karim, L.; Charloteaux, B.; Jamart, D.; Florin, A.; Lambert, C.; Rorive, A.; Jerusalem, G.; Leucci, E.; Dewaele, M.; Vooijs, M.; Leidel, S. A.; Georges, M.; Voz, M.; Peers, B.; Büttner, R.; Marine, J.; Chariot, A.; Close, P. Codon-specific translation reprogramming promotes resistance to targeted therapy. Nature 2018, 558, 605–609.

41. Delaunay, S.; Rapino, F.; Tharun, L.; Zhou, Z.; Heukamp, L.; Termathe, M.; Shostak, K.; Klevernic, I.; Florin, A.; Desmecht, H.; Desmet, C. J.; Nguyen, L.; Leidel, S. A.; Willis, A. E.; Büttner, R.; Chariot, A.; Close, P. Elp3 links tRNA modification to IRES- dependent translation of LEF1 to sustain metastasis in breast cancer. Journal of Experimental Medicine 2016, 213, 2503–2523.

42. Addis, L.; Ahn, J. W.; Dobson, R.; Dixit, A.; Ogilvie, C. M.; Pinto, D.; Vaags, A. K.; Coon, H.; Chaste, P.; Wilson, S.; Parr, J. R.; Andrieux, J.; Lenne, B.; Tumer, Z.; Leuzzi, V.; Aubell, K.; Koillinen, H.; Curran, S.; Marshall, C. R.; Scherer, S. W.; Strug, L. J.; Collier, D. A.; Pal, D. K. Microdeletions of ELP4 Are Associated with Language Impairment, Autism Spectrum Disorder, and Mental Retardation. Hum. Mutat. 2015, 36, 842–850.

43. Dogan, M.; Teralı, K.; Eroz, R.; Demirci, H.; Kocabay, K. Clinical and molecular findings in a Turkish family with an ultra-rare condition, ELP2-related neurodevelopmental disorder. Mol Biol Rep 2021, 48, 701–708.

44. Guo, W.; Russo, S.; Tuorto, F. Lost in translation: How neurons cope with tRNA decoding. BioEssays 2024, 2400107.

45. Anderson, S. J.; Coli, R.; Daly, I. W.; Kichula, E. A.; Rork, M. J.; Volpi, S. A.; Ekstein, J.; Rubin, B. Y. Familial dysautonomia is caused by mutations of the IKAP gene. American Journal of Human Genetics 2001, 68, 753–758.

46. Simpson, C. L.; Lemmens, R.; Miskiewicz, K.; Broom, W. J.; Hansen, V. K.; van Vught, Paul W J; Landers, J. E.; Sapp, P.; Van Den Bosch, L.; Knight, J.; Neale, B. M.; Turner, M. R.; Veldink, J. H.; Ophoff, R. A.; Tripathi, V. B.; Beleza, A.; Shah, M. N.; Proitsi, P.; Van Hoecke, A.; Carmeliet, P.; Horvitz, H. R.; Leigh, P. N.; Shaw, C. E.; van den Berg, Leonard H; Sham, P. C.; Powell, J. F.; Verstreken, P.; Brown, R. H., Jr; Robberecht, W.; Al-Chalabi, A. Variants of the elongator protein 3 (ELP3) gene are associated with motor neuron degeneration. Human Molecular Genetics 2009, 18, 472–481.

47. Bento-Abreu, A.; Jager, G.; Swinnen, B.; Rué, L.; Hendrickx, S.; Jones, A.; Staats, K. A.; Taes, I.; Eykens, C.; Nonneman, A.; Nuyts, R.; Timmers, M.; Silva, L.; Chariot, A.; Nguyen, L.; Ravits, J.; Lemmens, R.; Cabooter, D.; Van Den Bosch, L.; Van Damme, P.; Al-Chalabi, A.; Bystrom, A.; Robberecht, W. Elongator subunit 3 (ELP3) modifies ALS through tRNA modification. Human Molecular Genetics 2018, 27, 1276–1289.

48. Chovancova, E.; Pavelka, A.; Benes, P.; Strnad, O.; Brezovsky, J.; Kozlikova, B.; Gora, A.; Sustr, V.; Klavana, M.; Medek, P.; Biedermannova, L.; Sochor, J.; Damborsky, J. CAVER 3.0: A Tool for the Analysis of Transport Pathways in Dynamic Protein Structures. PLOS Computational Biology 2012, 8, e1002708.

49. Llewellyn, D. R.; O’Connor, C. Tracer studies of carboxylic acids. Part I. Acetic and pivalic acid. Journal of the Chemical Society 1964, 545–549.

50. 50. Samuel, D.; Silver, B. L. Oxygen Isotope Exchange Reactions of Organic Compounds. In Advances in Physical Organic Chemistry; Gold, V., Ed.; Academic Press: New York, 1965; pp 123–186.

51. Vavra, O.; Filipovic, J.; Plhak, J.; Bednar, D.; Marques, S. M.; Brezovsky, J.; Stourac, J.; Matyska, L.; Damborsky, J. CaverDock: a molecular docking-based tool to analyse ligand transport through protein tunnels and channels. Bioinformatics 2019, 35, 4986–4993.

52. Murail, S.; de Vries, S. J.; Rey, J.; Moroy, G.; Tufféry, P. SeamDock: An Interactive and Collaborative Online Docking Resource to Assist Small Compound Molecular Docking. Frontiers in Molecular Biosciences 2021, 8, 716466.

53. Tan, X.; Loke, H.; Fitch, S.; Lindahl, P. A. The Tunnel of Acetyl-Coenzyme A Synthase/Carbon Monoxide Dehydrogenase Regulates Delivery of CO to the Active Site. J. Am. Chem. Soc. 2005, 127, 5833–5839.

54. Biedermannová, L.; Prokop, Z.; Gora, A.; Chovancová, E.; Kovács, M.; Damborský, J.; Wade, R. C. A Single Mutation in a Tunnel to the Active Site Changes the Mechanism and Kinetics of Product Release in Haloalkane Dehalogenase LinB. Journal of Biological Chemistry 2012, 287, 29062–29074.

55. Brezovsky, J.; Babkova, P.; Degtjarik, O.; Fortova, A.; Gora, A.; Iermak, I.; Rezacova, P.; Dvorak, P.; Smatanova, I. K.; Prokop, Z.; Chaloupkova, R.; Damborsky, J. Engineering a de Novo Transport Tunnel. ACS Catal. 2016, 6, 7597–7610.

56. Farrugia, M. A.; Wang, B.; Feig, M.; Hausinger, R. P. Mutational and Computational Evidence That a Nickel-Transfer Tunnel in UreD Is Used for Activation of Klebsiella aerogenes Urease. Biochem 2015, 54, 6392–6401.

57. Lewis-Ballester, A.; Karkashon, S.; Batabyal, D.; Poulos, T. L.; Yeh, S. Inhibition Mechanisms of Human Indoleamine 2,3 Dioxygenase 1. J. Am. Chem. Soc. 2018, 140, 8518–8525.

58. Kokkonen, P.; Bednar, D.; Pinto, G.; Prokop, Z.; Damborsky, J. Engineering enzyme access tunnels. Biotechnology Advances 2019, 37, 107386.

59. Windsor, P.; Ouyang, H.; G. Da Costa, Joseph A; Rama Damodaran, A.; Chen, Y.; Bhagi-Damodaran, A. Gas Tunnel Engineering of Prolyl Hydroxylase Reprograms Hypoxia Signaling in Cells. Angew Chem Int Ed 2024, 63, e202409234.

60. Raushel, F. M.; Thoden, J. B.; Holden, H. M. Enzymes with Molecular Tunnels. Accounts of Chemical Research 2003, 36, 539–548.

61. Weeks, A.; Lund, L.; Raushel, F. M. Tunneling of intermediates in enzyme-catalyzed reactions. Current Opinion in Chemical Biology 2006, 10, 465–472.

62. Nicolet, Y.; Pagnier, A.; Zeppieri, L.; Martin, L.; Amara, P.; Fontecilla-camps, J. C. Crystal Structure of HydG from Carboxydothermus hydrogenoformans: A Trifunctional [FeFe]-Hydrogenase Maturase. ChemBioChem 2015, 16, 397–402.

63. Hyde, C. C.; Ahmedl, S. A.; Padlan, E. A.; Miles, E. W.; Davies, D. R. Three- dimensional Structure of the Tryptophan Synthase α2β2 Multienzyme Complex from Salmonella typhimurium. Journal of Biological Chemistry 1988, 263, 17857–17871.

64. Doukov, T. I.; Iverson, T. M.; Seravalli, J.; Ragsdale, S. W.; Drennan, C. L. A Ni-Fe- Cu Center in a Bifunctional Carbon Monoxide Dehydrogenase/ Acetyl-CoA Synthase. Science 2002, 298, 567–572.

65. Dinis, P.; Suess, D. L. M.; Fox, S. J.; Harmer, J. E.; Driesener, R. C.; De La Paz, L.; Swartz, J. R.; Essex, J. W.; Britt, R. D.; Roach, P. L. X-ray crystallographic and EPR spectroscopic analysis of HydG, a maturase in [FeFe]-hydrogenase H-cluster assembly. Proc. Natl. Acad. Sci. U. S. A. 2015, 112, 1362–1367.

66. Manjasetty, B. A.; Powlowski, J.; Vrielink, A. Crystal structure of a bifunctional aldolase-dehydrogenase: Sequestering a reactive and volatile intermediate. Proceedings of the National Academy of Sciences of the United States of America 2003, 100, 6992–6997.

67. Douangamath, A.; Walker, M.; Beismann-Driemeyer, S.; Cristina Vega-Fernandez, M.; Sterner, R.; Wilmanns, M. Structural Evidence for Ammonia Tunneling across the (βα)8 Barrel of the Imidazole Glycerol Phosphate Synthase Bienzyme Complex. Structure 2002, 10, 185–193.

68. Chaudhuri, B. N.; Lange, S. C.; Myers, R. S.; Davisson, V. J.; Smith, J. L. Toward Understanding the Mechanism of the Complex Cyclization Reaction Catalyzed by Imidazole Glycerolphosphate Synthase: Crystal Structures of a Ternary Complex and the Free Enzyme, Biochem 2003, 42, 7003–7012.

69. Oberg, N.; Precord, T. W.; Mitchell, D. A.; Gerlt, J. A. RadicalSAM.org: A Resource to Interpret Sequence-Function Space and Discover New Radical SAM Enzyme Chemistry. ACS Bio Med Chem Au 2021, 2, 22–35.

70. 70. Uniprot Consortium UniProt: the universal protein knowledgebase. Nucleic Acids Res 2018, 45, D158–D169.

71. Berman, H.; Henrick, K.; Nakamura, H. Announcing the worldwide Protein Data Bank. Nature Structural & Molecular Biology 2003, 10, 980.

72. 72. Holliday, G. L.; Akiva, E.; Meng, E. C.; Brown, S. D.; Calhoun, S.; Pieper, U.; Sali, A.; Booker, S. J.; Babbitt, P. C. Atlas of the Radical SAM Superfamily: Divergent Evolution of Function Using a “Plug & Play” Domain. In Methods in Enzymology; Bandarian, V., Ed.; Academic Press: 2018; Vol. 606, pp 1–71.

73. Shibata, N.; Toraya, T. Molecular architectures and functions of radical enzymes and their (re)activating proteins. J Biochem 2015, 158, 271–292.

74. Chan, P. P.; Lowe, T. M. GtRNAdb 2.0: an expanded database of transfer RNA genes identified in complete and draft genomes. Nucleic Acids Res 2015, 44, D184–D189.

75. Chramiec-Głąbik, A.; Rawski, M.; Glatt, S.; Lin, T. Electrophoretic Mobility Shift Assay (EMSA) and Microscale Thermophoresis (MST) Methods to Measure Interactions Between tRNAs and Their Modifying Enzymes. In *RNA-Protein Complexes and Interactions: Methods and Protocols*; Lin, R., Ed.; Humana: New York, NY, 2023; pp 29–53.

76. Schneider, C. A.; Rasband, W. S.; Eliceiri, K. W. NIH Image to ImageJ: 25 years of image analysis. Nature Methods 2012, 9, 671–675.

