## Supporting Information for "Elp3 uses a conserved molecular tunnel to transport acetate between distant active sites and catalyze tRNA wobble base modification"

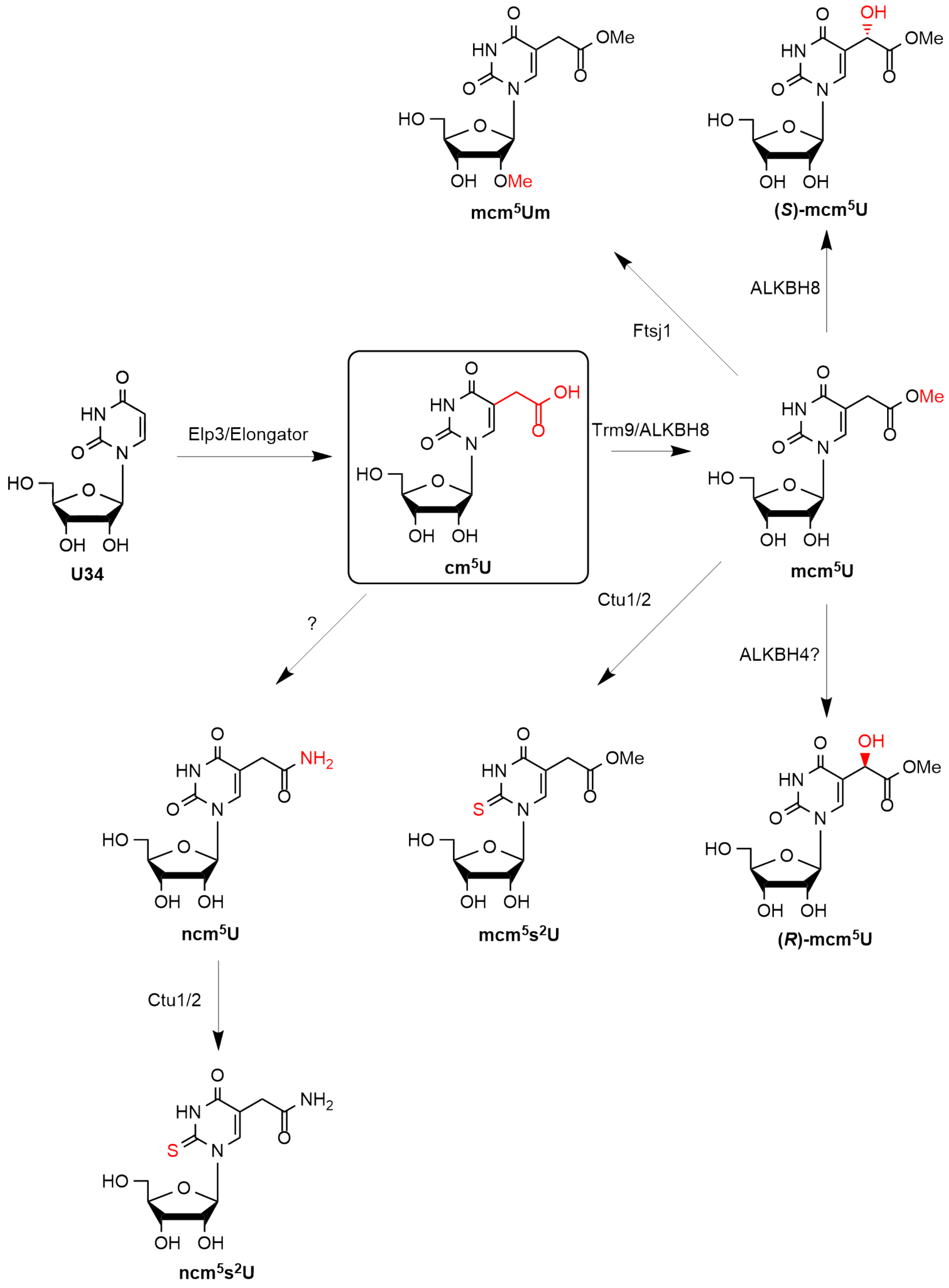


**Figure S1. 5-carboxymethyluridine (cm$^5$U)-derived eukaryotic tRNA modifications and modification enzymes.** Elp3 and Elongator install the central intermediate tRNA modification cm$^5$U at the wobble base position (U34). cm$^5$U is further elaborated by additional tRNA modification enzymes to produce a family of cm$^5$U-derived modifications that impact translation efficiency and fidelity. Eukaryotic cm$^5$U-derived modifications and their known or speculated tRNA modification enzymes are shown.

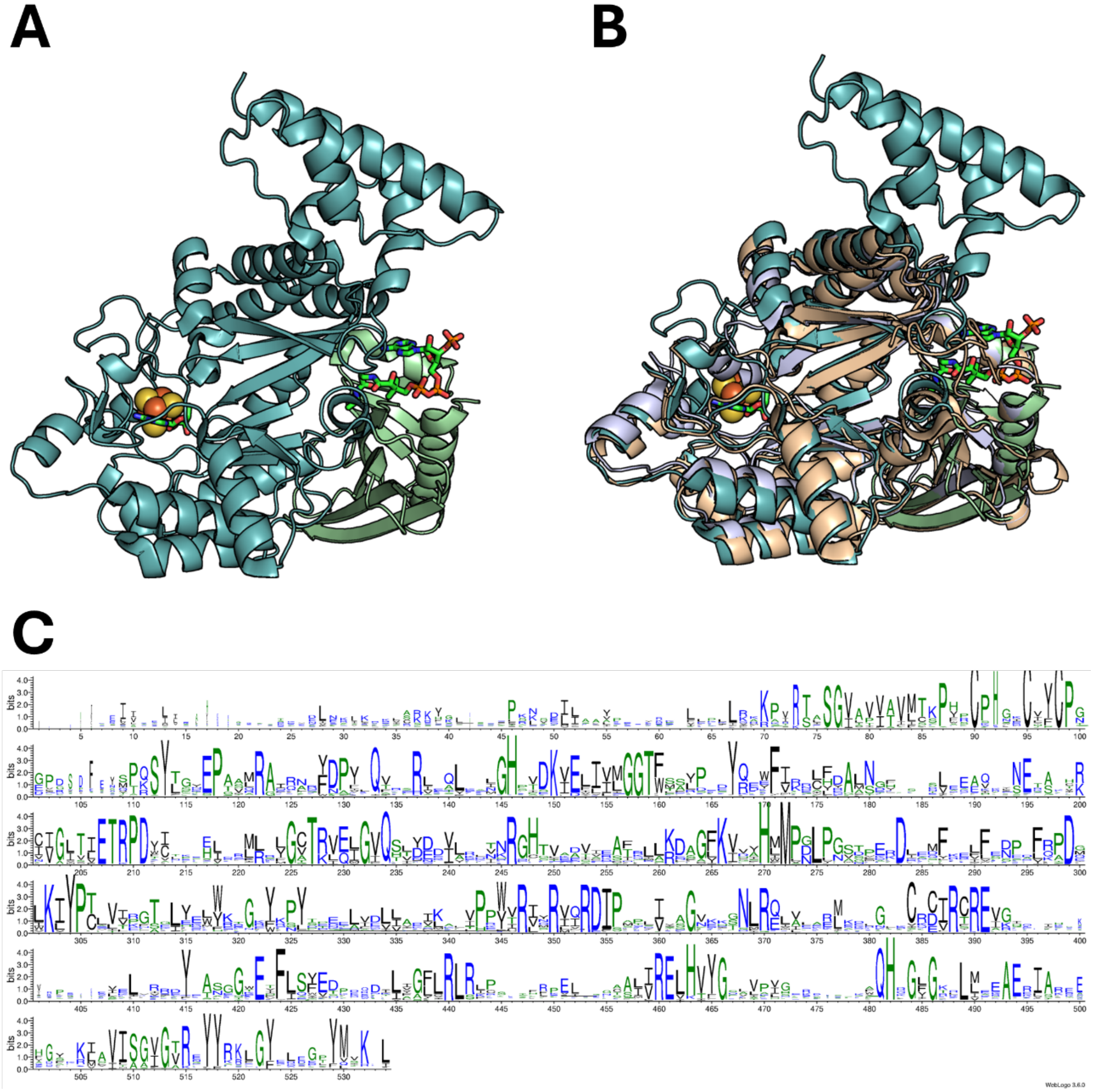


**Figure S2. Elp3 has a highly conserved sequence and tertiary structure. (A)** Structure of *Saccharomyces cerevisiae* Elp3 (rSAM domain in teal and KAT domain in light green, PDB 8ASW) with 5′-dA and AcCoA analogue desulfo-CoA (aligned from PDB 6IA6) shown as green sticks. **(B)** Elp3 structural alignment from *S. cerevisiae* (eukaryote, rSAM domain in teal and KAT domain in light green, PDB 8ASW), *Dehalococcoides mccartyi* (bacteria, light blue, PDB 6IA6), and *Methanocaldococcus infernus* (*Min*, archaea, tan, PDB 6IA8). 5′-dA (PDB 8ASW) and AcCoA analog desulfo-CoA (PDB 6IA6) are shown as green sticks and the [4Fe-4S] cluster (PDB 8ASW) is shown in orange and yellow spheres. **(C)** Elp3 sequence alignment using DeepMSA2[1] with an alignment depth (Nf) of 18.03 from 568 sequences spanning archaea, bacteria, and both lower and higher eukaryotes. Visualized with Weblogo.[2] Residues are numbered according to the *Min* sequence.

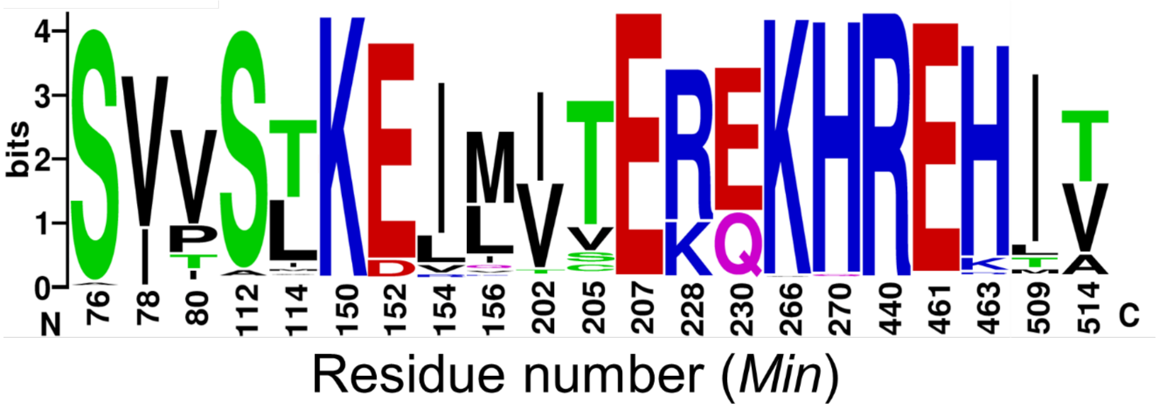


**Figure S3. The residues forming Elp3's molecular tunnel are highly conserved.** Tunnel-lining residues are shown and numbered according to the *Min* sequence. Alignment generated with DeepMSA2[1] (full alignment shown in Figure S2, Nf = 18.03, 568 sequences spanning archaea, bacteria, and eukaryotes) and visualized with Weblogo.[2]

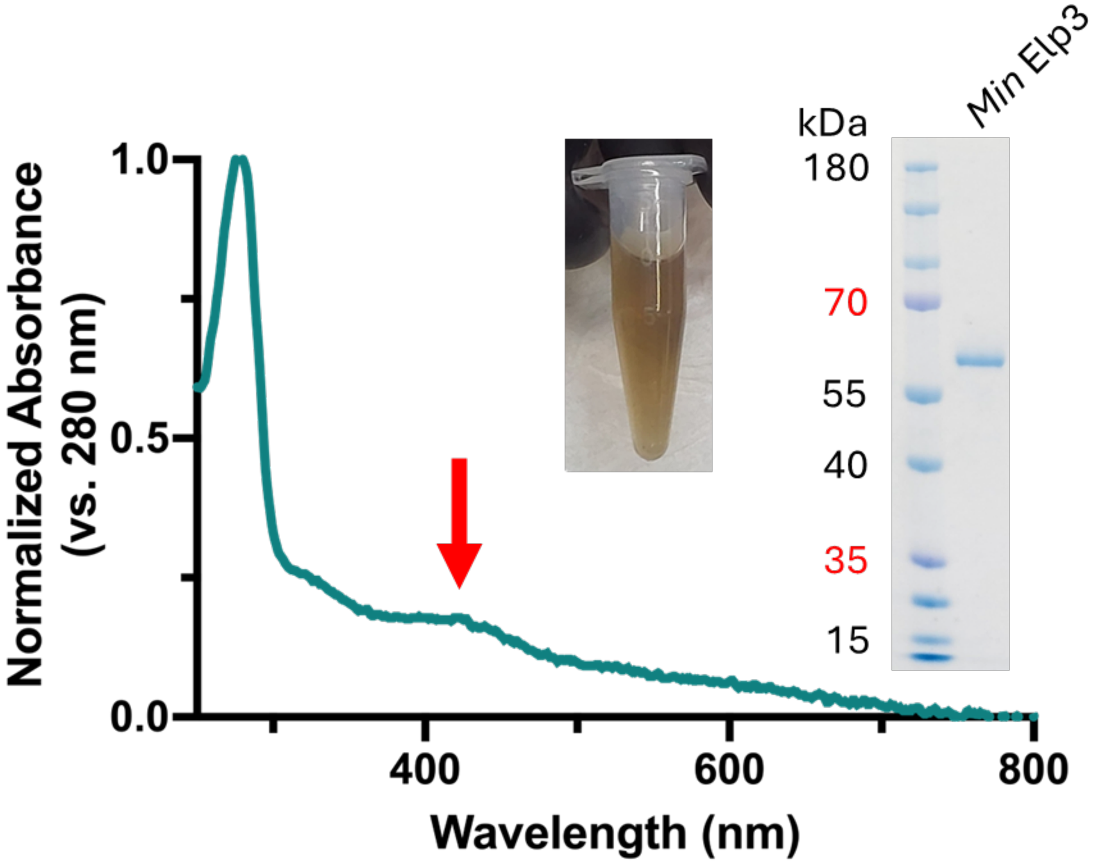


**Figure S4. Characterization of WT *Min* Elp3.** The UV-Vis spectrum of purified, reconstituted *Min* Elp3 has a shoulder at 420 nm (red arrow) consistent with the presence of a [4Fe-4S] cluster. The inset images show the characteristic brown-yellow color of *Min* Elp3 following reconstitution and its purity as assessed by gel electrophoresis. The *Min* Elp3 construct is predicted to have a mass of 65.0 kDa.

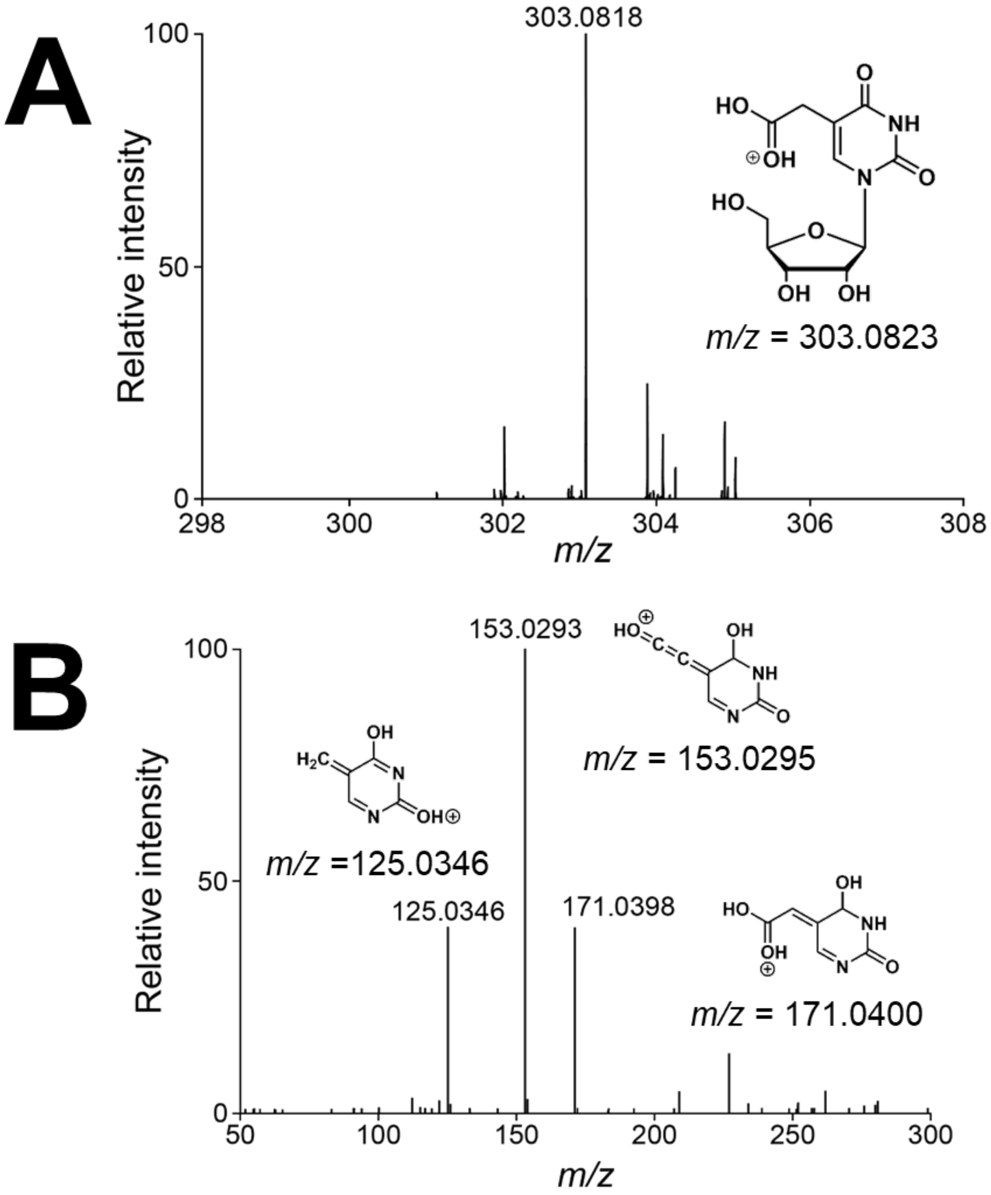


**Figure S5. Mass spectra of $cm^5U$ product from Elp3 reactions. (A)** LC-MS analysis of nuclease-digested tRNA nucleosides confirms that Elp3 reactions with non-isotopically labeled AcCoA produce $cm^5U$. **(B)** MS/MS analysis of the $cm^5U$ nucleoside product peak revealed characteristic mass fragments of $cm^5U$ (compared to a commercial $cm^5U$ standard). The tRNA modification reaction was performed with 5 µM Elp3, 4.4 µM tRNA, 25 µM SAM, 0.5 mM dithionite, and 27.5 µM AcCoA. MS ion structures and masses were predicted with CFM-ID 4.0.[3]

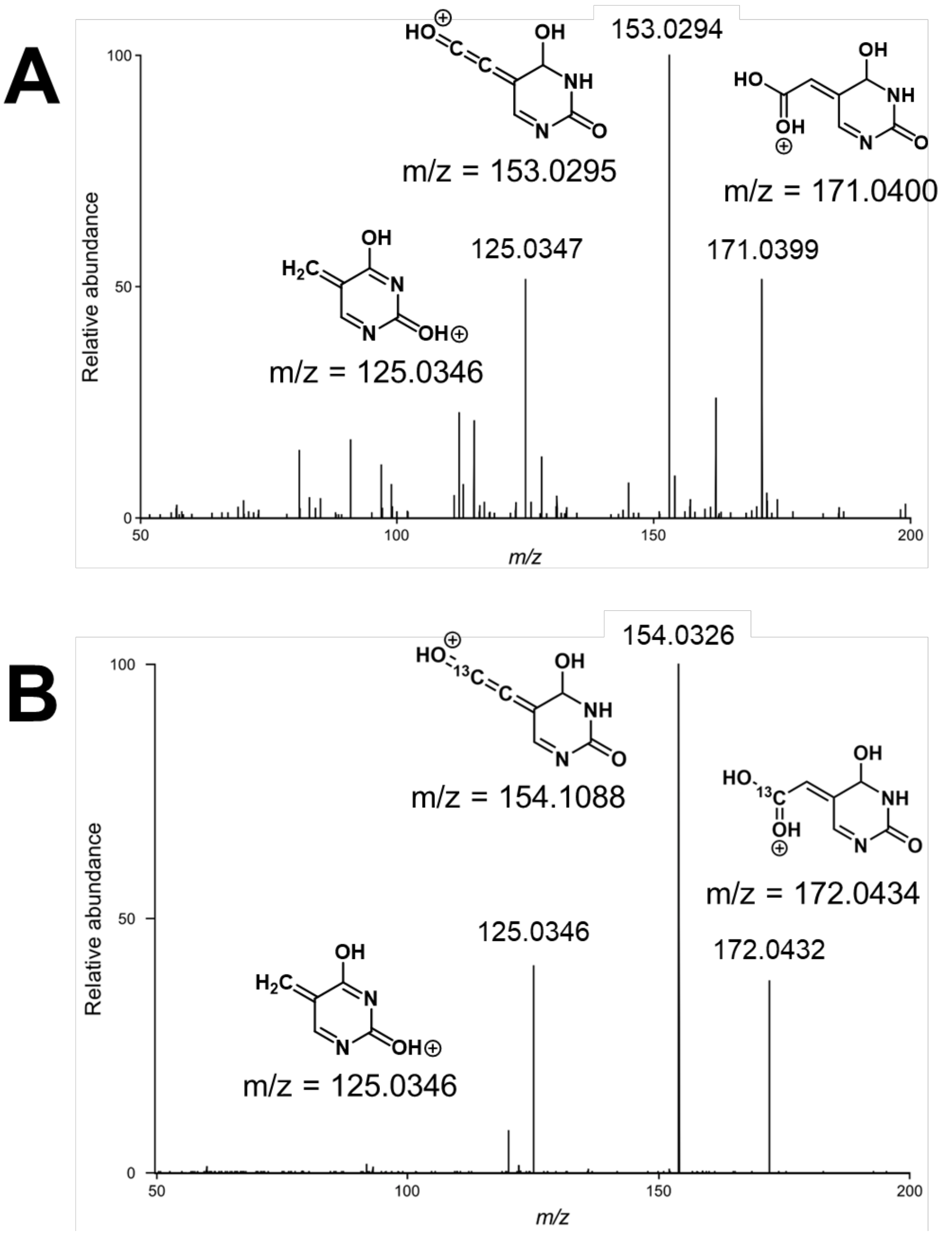


**Figure S6. MS/MS spectra of cm⁵U product from (A) $^{12}$C and (B) $^{13}$C acetate reactions. (A)** MS/MS analysis of the $^{12}$C (non-isotopically labeled) acetate reaction cm$^5$U nucleoside product peak revealed characteristic mass fragments of cm$^5$U. **(B)** MS/MS analysis of the $^{13}$C acetate reaction cm$^5$U nucleoside product peak revealed characteristic mass fragments of cm$^5$U with appropriate +1 Da mass shifts based on the location of the $^{13}$C label. tRNA modification reactions were performed with 5 µM Elp3, 4.4 µM tRNA, 25 µM SAM, 0.5 mM dithionite, and 10 mM $^{12}$C or $^{13}$C acetate. MS ion structures and masses were predicted with CFM-ID 4.0.[3]

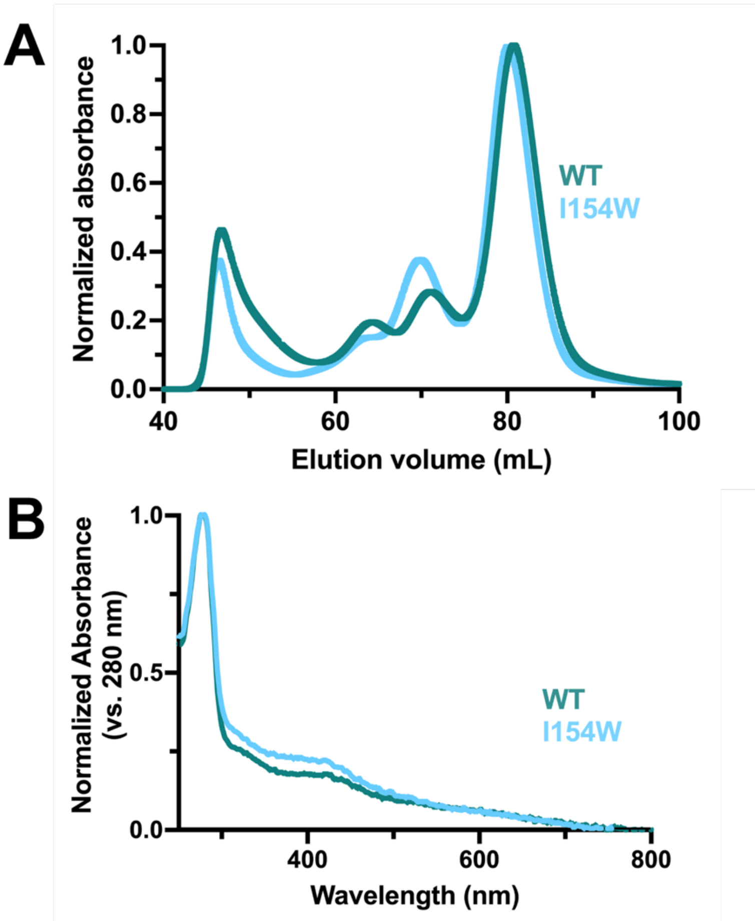


**Figure S7. Comparison of purified and reconstituted I154W vs WT *Min* Elp3. (A)** Normalized size exclusion chromatography (SEC) traces of WT (teal) and I154W (light blue) *Min* Elp3. **(B)** UV-Vis spectra of WT (teal) and I154W (light blue) *Min* Elp3 after reconstitution; both spectra show similar, characteristic [4Fe-4S] absorbance peaks at 420 nm.

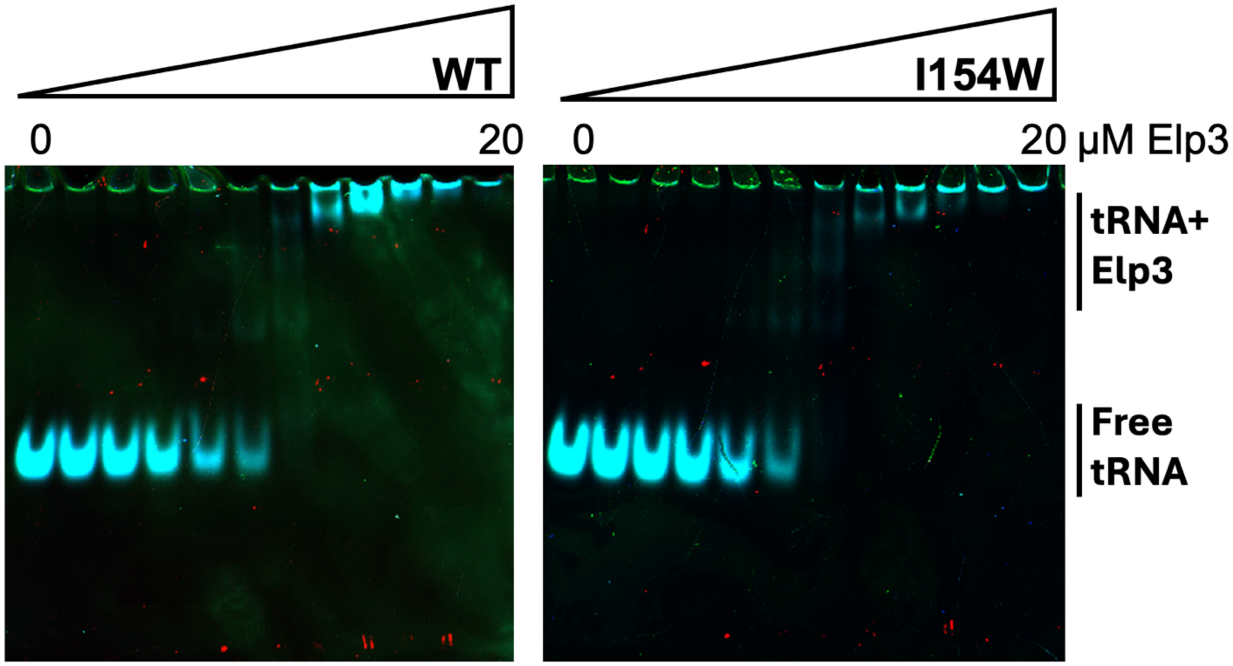


**Figure S8. Representative EMSA gels used to measure tRNA binding to Elp3.** 250 nM tRNA was incubated with 0 – 20 µM Elp3, free and bound tRNA species were separated on a 5% TBE gel, and tRNA was visualized with SYBR gold staining (stained tRNA is cyan). Elp3-tRNA complexes migrate very slowly on the gel and were difficult to reliably quantify, so fraction bound was calculated by quantifying the amount of free tRNA at each Elp3 concentration compared to total tRNA (0 µM Elp3).

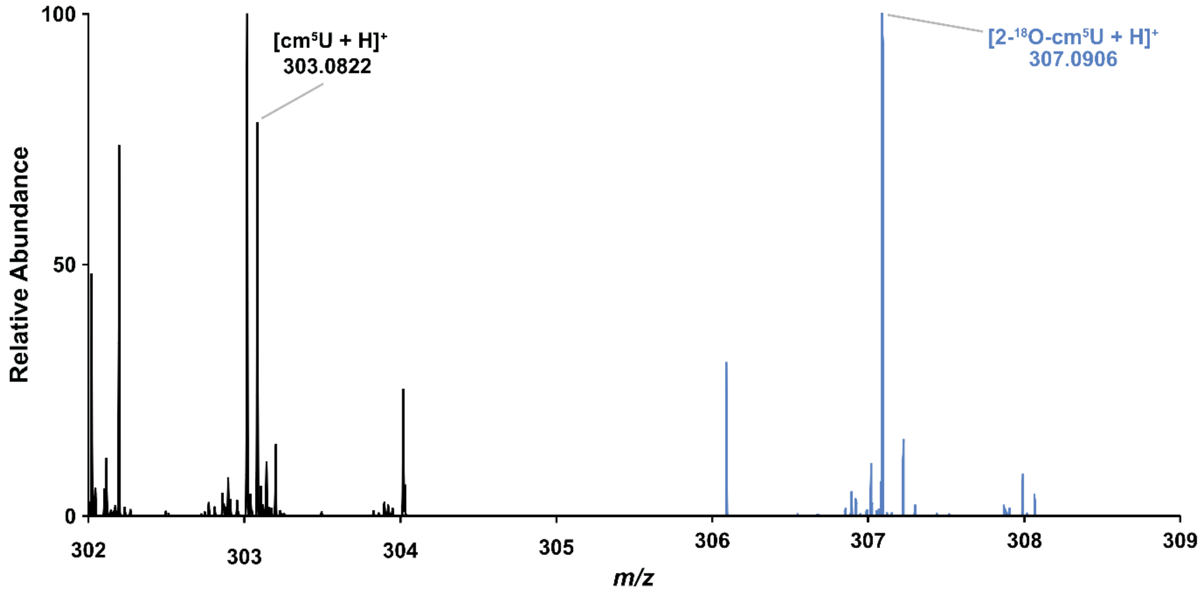


**Figure S9. Uncropped mass spectra of the $^{12}C$ and $^{18}O_2$ acetate cm$^5$U nucleoside products.** The black LC-MS spectrum shows the cm$^5$U product from $^{12}C$-acetate reactions with Elp3; the blue LC-MS spectrum shows the cm$^5$U product from $^{18}O_2$-acetate reactions with Elp3, showing the +4 Da shift from the isotopically unlabeled acetate reactions. This is the same data as shown in Figure 3B, but with an uncropped x-axis.

**A**

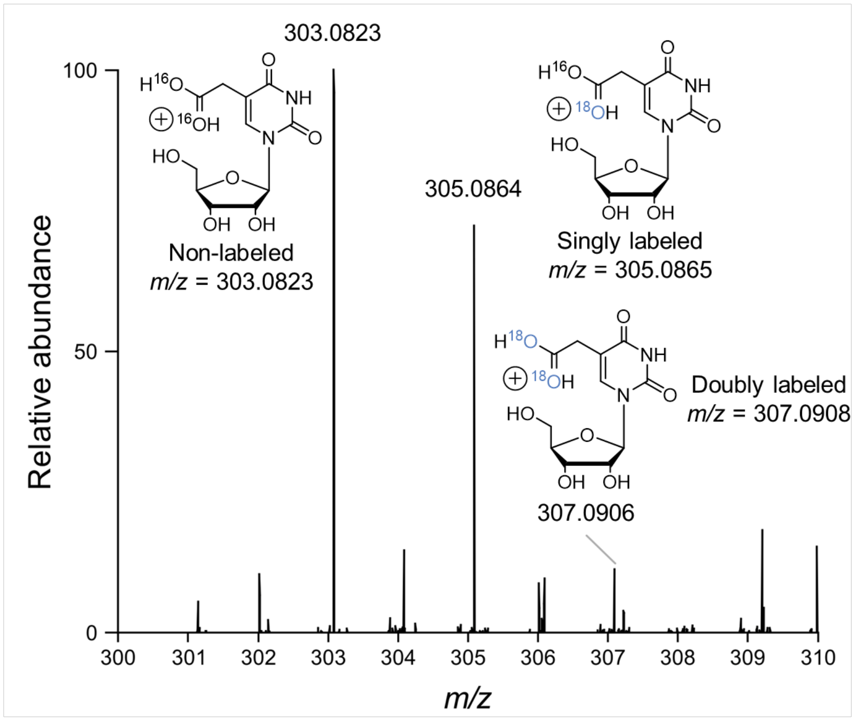


**B**

**(i)** **Isotope ratio relative to $^{16}O$ cm$^5$U**

| Ratio | 0-$^{18}O$ (all $^{16}O$) | 1-$^{18}O$ (+2 Da) | 2-$^{18}O$ (+4 Da) |
|---|---|---|---|
| Theoretical | 99.91 | 1.71 | 0.01 |
| Observed | 100 | 71.47 | 10.44 |

**(ii)** **Isotope ratio relative to singly-labeled $^{18}O$ cm$^5$U**

| Ratio | 1-$^{18}O$ (+2 Da) | 2-$^{18}O$ (+4 Da) |
|---|---|---|
| Theoretical | 99.97 | 1.51 |
| Observed | 100 | 11.70 |

**Figure S10. $^{18}O$ ratio analysis for Elp3-mediated reactions with $^{18}O_2$-acetate. (A)** LC-MS analysis of the *in vitro* activity assay with $^{18}O_2$ acetate showing doubly and singly $^{18}O$-labeled cm$^5$U, as well as nonlabelled cm$^5$U. The cm$^5$U [M+1] masses were predicted with enviPat.[4] **(B)** An analysis of the oxygen isotope ratios between different isotopically labeled cm$^5$U products reveals that doubly $^{18}O$-labeled cm$^5$U is observed at a relative abundance **(i)** ~1000-fold higher than would be expected in normal abundance (theoretical) cm$^5$U and **(ii)** ~8-fold higher than would be expected in a singly $^{18}O$-labeled-only cm$^5$U sample. Theoretical ratios expected for the specific LC-MS instrument used were predicted with enviPat (resolution setting: Q-Exactive, ExactivePlus_R140000@200`) and observed ratios were calculated using the peak areas of the various cm$^5$U isotopes.

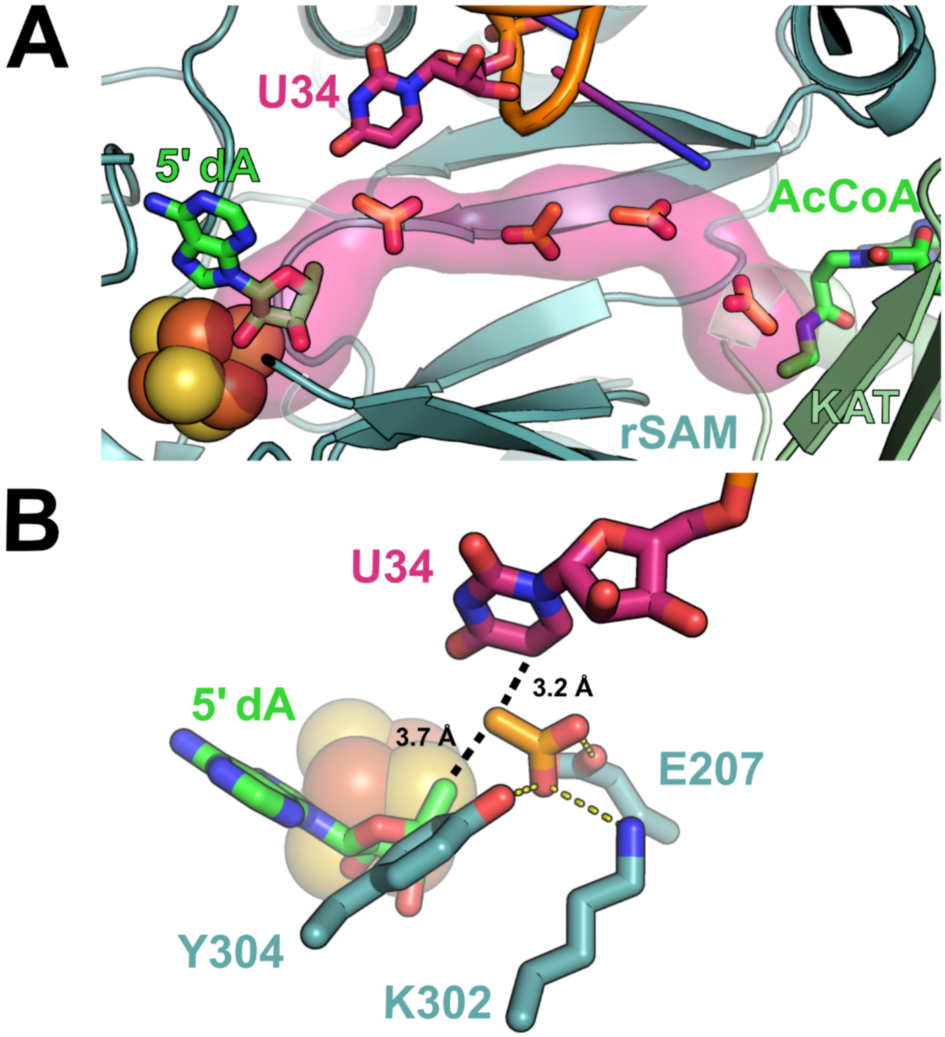


**Figure S11. Molecular docking provides hypotheses about how acetate may occupy the Elp3 molecular tunnel. (A)** Docking acetate in the yeast structure of Elp3 bound to tRNA (PDB 8ASW) with CaverDock[5] suggests that acetate can reasonably be accommodated at many positions along the molecular tunnel. The Elp3 rSAM domain is colored teal, the Elp3 KAT domain is colored light green, the calculated tunnel is shown in pink, and 4 CaverDock-predicted acetate poses are shown in orange sticks along the length of the tunnel; 5′-dA and AcCoA analog desulfo-CoA (aligned from PDB 6IA6) are shown in green sticks; tRNA and substrate tRNA base U34 are shown in orange and pink, respectively. **(B)** CaverDock modeling of acetate in the rSAM active site. The CaverDock pose shown here provides a similar, but alternative, conformation of acetate in the Elp3 active site, as compared to the SeamDock model shown in Figure 4A. Both this model and the one in Figure 4A show how acetate may be oriented by noncovalent interactions (yellow dotted lines) in the rSAM active site that position the acetate methyl group at a reasonable distance for sequential reaction with 5′-dA· and tRNA U34.

| *Min* | Yeast | Human |
|---|---|---|
| E207 | E230 | E221 |
| K302 | K325 | K316 |
| Y304 | Y327 | Y318 |

**Table S1. Numbering of key, conserved rSAM active site residues for archaeal *Methanocaldococcus infernus* (*Min*), yeast, and human Elp3.**

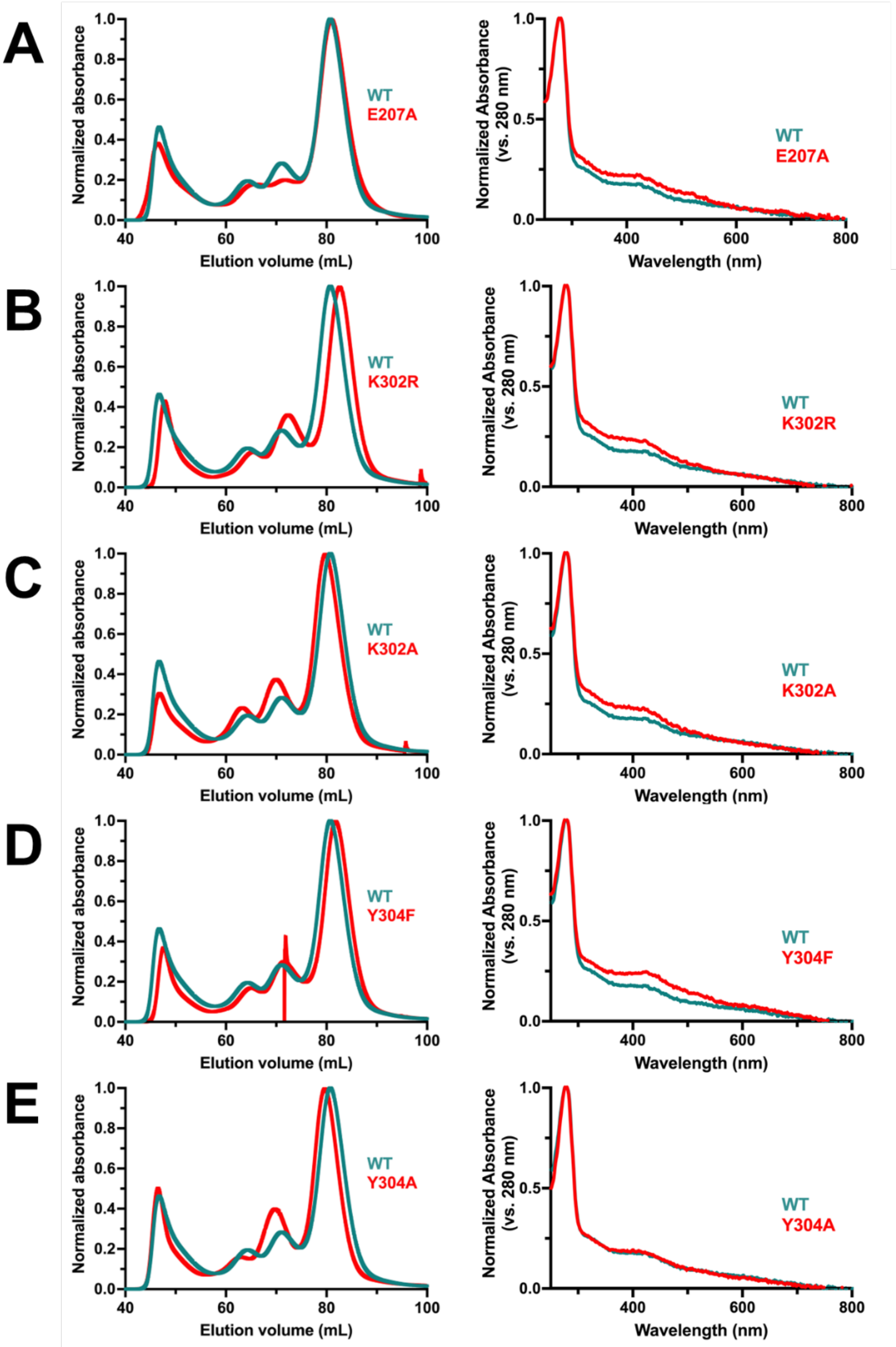


**Figure S12. Comparison of the purified and reconstituted rSAM active site variants to WT *Min* Elp3.** SEC traces (left) and UV-Vis spectra (right) of the rSAM site variants E207A **(A)**, K302R **(B)**, K302A **(C)**, Y304F **(D)**, and Y304A **(E)** compared to WT Elp3. All Elp3 mutants show similar SEC traces and similar, characteristic [4Fe-4S] absorbance peaks at 420 nm, as compared to WT *Min* Elp3.

### References


1. Zheng, W.; Wuyun, Q.; Li, Y.; Zhang, C.; Freddolino, P. L.; Zhang, Y. Improving deep learning protein monomer and complex structure prediction using DeepMSA2 with huge metagenomics data. *Nat Methods* **2024***, 21*, 279–289.

2. Crooks, G. E.; Hon, G.; Chandonia, J.; Brenner, S. E. WebLogo: A Sequence Logo Generator. *Genome Res.* **2004***, 14*, 1188-1190.

3. Wang, F.; Allen, D.; Tian, S.; Oler, E.; Gautam, V.; Greiner, R.; Metz, T. O.; Wishart, D. S. CFM-ID 4.0 – a web server for accurate MS-based metabolite identification. *Nucleic Acids Research* **2022***, 50*, W165–W174.

4. Loos, M.; Gerber, C.; Corona, F.; Hollender, J.; Singer, H. Accelerated Isotope Fine Structure Calculation Using Pruned Transition Trees. *Anal. Chem.* **2015***, 87*, 5738-5744.

5. Vavra, O.; Filipovic, J.; Plhak, J.; Bednar, D.; Marques, S. M.; Brezovsky, J.; Stourac, J.; Matyska, L.; Damborsky, J. CaverDock: a molecular docking-based tool to analyse ligand transport through protein tunnels and channels. *Bioinformatics* **2019***, 35*, 4986–4993.
